# Molecular basis of the urate transporter URAT1 inhibition by gout drugs

**DOI:** 10.1101/2024.09.11.612563

**Authors:** Yang Suo, Justin G. Fedor, Han Zhang, Kalina Tsolova, Xiaoyu Shi, Kedar Sharma, Shweta Kumari, Mario Borgnia, Peng Zhan, Wonpil Im, Seok-Yong Lee

## Abstract

Hyperuricemia is a condition when uric acid, a waste product of purine metabolism, accumulates in the blood^1^. Untreated hyperuricemia can lead to crystal formation of monosodium urate in the joints, causing a painful inflammatory disease known as gout. These conditions are associated with many other diseases and affect a significant and increasing proportion of the population^2–4^. The human urate transporter 1 (URAT1) is responsible for the reabsorption of ∼90% of uric acid in the kidneys back into the blood, making it a primary target for treating hyperuricemia and gout^5^. Despite decades of research and development, clinically available URAT1 inhibitors have limitations because the molecular basis of URAT1 inhibition by gout drugs remains unknown^5^. Here we present cryo-electron microscopy structures of URAT1 alone and in complex with three clinically relevant inhibitors: benzbromarone, lesinurad, and the novel compound TD-3. Together with functional experiments and molecular dynamics simulations, we reveal that these inhibitors bind selectively to URAT1 in inward-open states. Furthermore, we discover differences in the inhibitor dependent URAT1 conformations as well as interaction networks, which contribute to drug specificity. Our findings illuminate a general theme for URAT1 inhibition, paving the way for the design of next-generation URAT1 inhibitors in the treatment of gout and hyperuricemia.

## Main

Gout is a disease that afflicts up to 6.8% of the population globally^2^ and is the most common form of inflammatory arthritis, particularly among men in developed countries^2,3^. Characterized by recurrent episodes of acute inflammatory arthritis, gout is primarily driven by the deposition of monosodium urate (MSU) crystals within joints. Hyperuricemia, a major risk factor for gout, is characterized by an accumulation of uric acid in the blood and is increasingly recognized as a potential contributor to a spectrum of comorbidities including cardiovascular diseases, renal disorders, kidney failure, diabetes, and metabolic syndrome^4,6–10^. Currently, 21% of Americans are diagnosed with hyperuricemia^4^ and global prevalence is estimated up to 36% in different populations^6^. Despite these implications, the management of hyperuricemia and gout remains suboptimal, largely due to limitations in current pharmacological interventions. Unfortunately, the number of gout cases is rapidly surging, with the prevalence of gout increasing globally from 1990 to 2019 by ∼21%, and by 90.6% for men in the United States^3^. This not only bears considerable impact on individual quality of life, but a quickly growing burden for public health. Much needed improvements in treatments are therefore needed through a better understanding of the causes of gout and the pharmacological targets.

The human urate transporter 1 (URAT1) is encoded by the *SLC22A12* gene which belongs to the SLC22 family of organic cation/anion transporters. URAT1, primarily expressed on the luminal side of the renal proximal tubule, uptakes urate in exchange for exporting monocarboxylates^11^, serving as a specific and major regulator of uric acid reabsorption from the urine (Fig 1a)^11,12^. Approximately 90% of the urate filtered from glomeruli is reabsorbed back to the bloodstream, while only 10% is excreted in the urine^1^. Reabsorption of urate is largely mediated by URAT1, making it the critical target for the treatment of hyperuricemia and gout^1,12,13^. Case in point, 90% of hypouricemia cases are linked to nonfunctional mutations in URAT1, where the vast majority of mutations are protective against gout and hyperuricemia^5^. Inhibition of URAT1 is therefore an effective strategy to promote uric acid excretion to mitigate the risk of hyperuricemia-related complications, including gout^1^.

**Figure 1.**
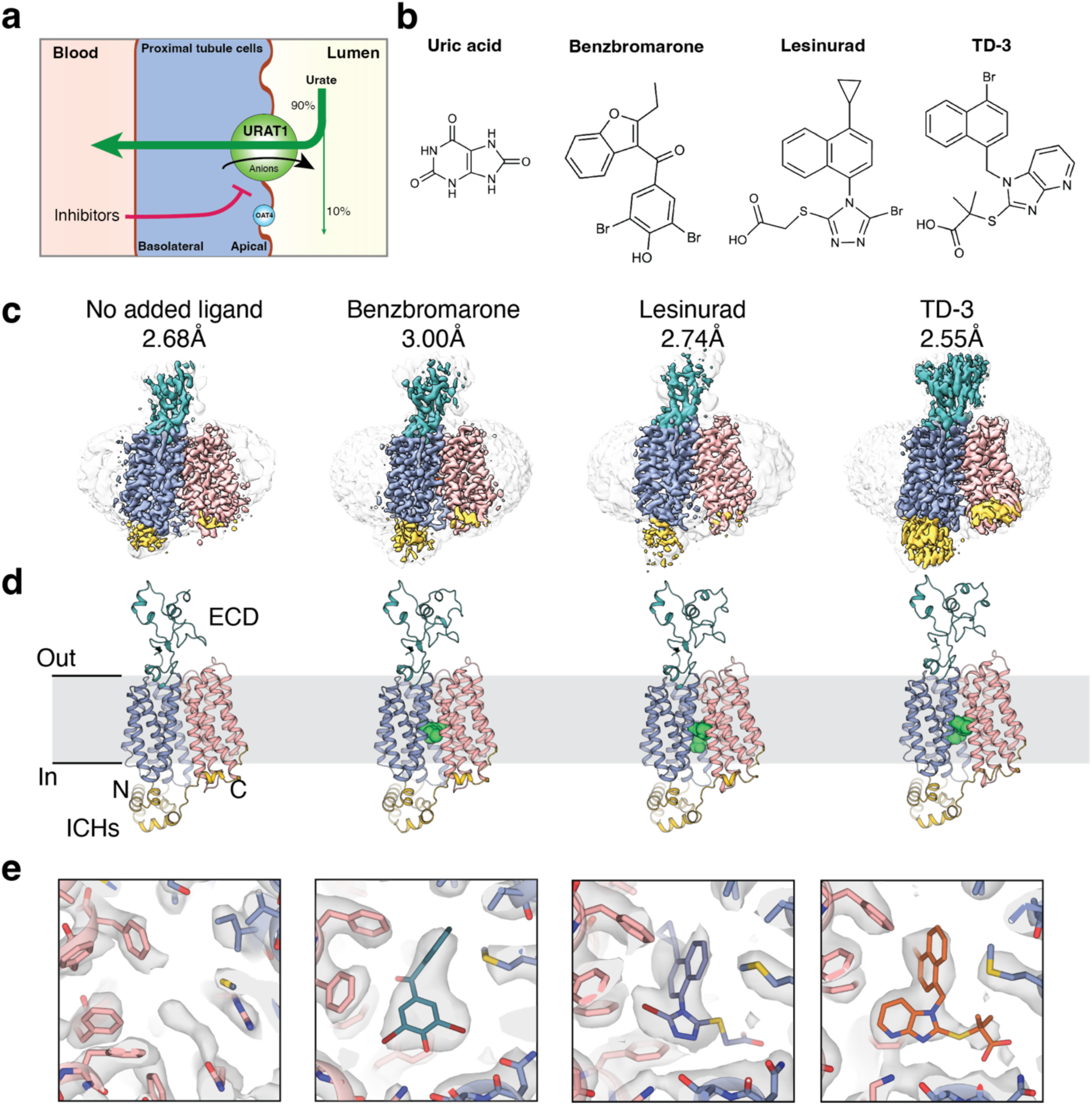
URAT1 biology and structure. **a,** The role of URAT1 in urate reabsorption in the kidney proximal tubule epithelium. **b,** Chemical structures of URAT1 substrate and inhibitors. **c-e,** cryo-EM reconstructions, structures, and map of the central binding cavity for URAT1_CS_ alone and in complex with benzbromarone (BBR-URAT1_CS_), lesinurad (LESU-URAT1_CS_) and TD-3 (TD-3-URAT1_CS_).

Despite the clear therapeutic potential of targeting URAT1, the development of specific and potent inhibitors has proven challenging. Benzbromarone (BBR) has been used to treat gout for more than 30 years^14^. Although it is a potent inhibitor of URAT1 and effective at lowering serum uric acid concentrations, reports of hepatoxicity have led to reduction in its use^15,16^. In 2015 the FDA approved lesinurad (LESU) as a novel inhibitor of URAT1 for the treatment of gout and hyperuricemia, but it must be co-administered with the xanthine oxidase inhibitor allopurinol due to its toxicity and low efficacy^17^. More recently, utilizing lesinurad as a lead compound, a group of novel bicyclic imidazopyridines were developed as URAT1 inhibitors^18^. Among these, a compound named TD-3 (compound **23** in the original study) exhibits exceptional properties, including excellent ability to lower serum urate *in vivo*, favorable safety and pharmacokinetic properties, oral bioavailability, and potent *in vitro* inhibition (IC_50_ 1.36 µM), surpassing lesinurad in all aspects^18^. Overall, TD-3 shows promise as a drug candidate for hyperuricemia and gout^18^.

These issues and progresses highlight the pressing need for the development of more selective and safer URAT1 inhibitors. Therefore, we sought to better understand URAT1 inhibition through functional assays and cryo-EM with a focus on the inhibitors BBR, LESU and TD-3 with the aim of identifying key structural features of URAT1 that could be leveraged for future drug development.

### URAT1_CS_ binds inhibitors in the inward-open conformation

Wild-type human URAT1 exhibits poor expression and stability when expressed in HEK293S GnTI^-^ cells, hindering structural elucidation. We turned to consensus mutagenesis to improve protein yield and stability, as previously implemented in our laboratory^19^. We obtained a construct with 100% sequence identity to human URAT1 in the central ligand binding cavity, with an overall 91% sequence identity to human URAT1 (Extended Data Figure 1a, Supplemental Figure 1). We hereafter refer to this construct as URAT1_CS_, which shows superior expression yields and stability by size exclusion chromatography (Extended Data Figure 1b). However, [^14^C]-uric acid (UA) uptake assays in HEK293T cells transiently expressing hURAT1 and URAT1_CS_ show that URAT1_CS_ has substantially weaker uptake activity compared with hURAT1 (Extended Data Figure 1c). This suggests URAT1_CS_ adopts an over-stabilized conformation but is still capable of turnover. Importantly, measurement of the IC_50_ for TD-3 in HEK293T cells expressing hURAT1 versus URAT1_CS_ show that URAT1_CS_ binds TD-3 with a similar affinity compared to hURAT1 (Extended Data Figure 1e). So despite a very slow turnover, the inhibitor binding site and properties of the central cavity is preserved.

We determined the cryo-electron microscopy (cryo-EM) structures of URAT1_CS_ alone at 2.68 Å, in complex with benzbromarone (BBR-URAT1_CS_) at 3.00 Å, in complex with lesinurad (LESU-URAT1_CS_) at 2.74 Å and in complex with TD-3 (TD-3-URAT1_CS_) at 2.55 Å (Fig. 1c, 1d, Extended Data Figure 2,3, Table 1). Robust cryo-EM densities within the central cavity were identified, and the corresponding inhibitors were unambiguously modeled. There is also a weaker density in the central cavity of URAT1_CS_ alone, likely from an endogenous molecule, but its position and shape are distinct from those of the inhibitors (Fig. 1e, Extended Data Figure 4).

**Table 1.**
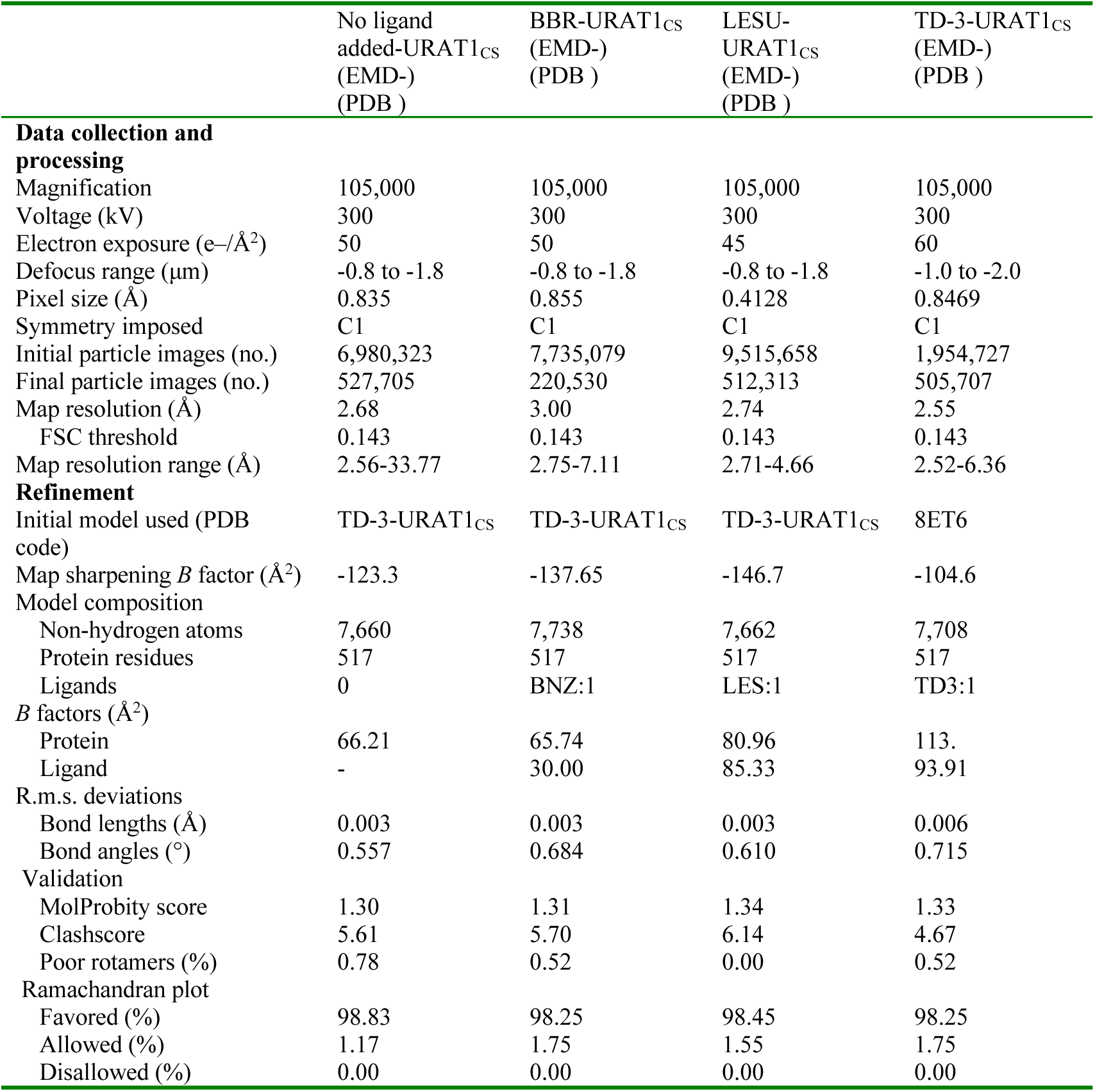
Cryo-EM data collection, refinement, and validation statistics.

|  | No ligand<br>added-URAT1<sub>CS</sub><br>(EMD-)<br>(PDB ) | BBR-URAT1<sub>CS</sub><br>(EMD-)<br>(PDB ) | LESU-URAT1<sub>CS</sub><br>(EMD-)<br>(PDB ) | TD-3-URAT1<sub>CS</sub><br>(EMD-)<br>(PDB ) |
| --- | --- | --- | --- | --- |
| **Data collection and processing** |  |  |  |  |
| Magnification | 105,000 | 105,000 | 105,000 | 105,000 |
| Voltage (kV) | 300 | 300 | 300 | 300 |
| Electron exposure (e-/Å<sup>2</sup>) | 50 | 50 | 45 | 60 |
| Defocus range (µm) | -0.8 to -1.8 | -0.8 to -1.8 | -0.8 to -1.8 | -1.0 to -2.0 |
| Pixel size (Å) | 0.835 | 0.855 | 0.4128 | 0.8469 |
| Symmetry imposed | C1 | C1 | C1 | C1 |
| Initial particle images (no.) | 6,980,323 | 7,735,079 | 9,515,658 | 1,954,727 |
| Final particle images (no.) | 527,705 | 220,530 | 512,313 | 505,707 |
| Map resolution (Å) | 2.68 | 3.00 | 2.74 | 2.55 |
| FSC threshold | 0.143 | 0.143 | 0.143 | 0.143 |
| Map resolution range (Å) | 2.56-33.77 | 2.75-7.11 | 2.71-4.66 | 2.52-6.36 |
| **Refinement** |  |  |  |  |
| Initial model used (PDB code) | TD-3-URAT1<sub>CS</sub> | TD-3-URAT1<sub>CS</sub> | TD-3-URAT1<sub>CS</sub> | 8ET6 |
| Map sharpening *B* factor (Å<sup>2</sup>) | -123.3 | -137.65 | -146.7 | -104.6 |
| **Model composition** |  |  |  |  |
| Non-hydrogen atoms | 7,660 | 7,738 | 7,662 | 7,708 |
| Protein residues | 517 | 517 | 517 | 517 |
| Ligands | 0 | BNZ:1 | LES:1 | TD3:1 |
| ***B* factors (Å<sup>2</sup>)** |  |  |  |  |
| Protein | 66.21 | 65.74 | 80.96 | 113. |
| Ligand | - | 30.00 | 85.33 | 93.91 |
| **R.m.s. deviations** |  |  |  |  |
| Bond lengths (Å) | 0.003 | 0.003 | 0.003 | 0.006 |
| Bond angles (°) | 0.557 | 0.684 | 0.610 | 0.715 |
| **Validation** |  |  |  |  |
| MolProbity score | 1.30 | 1.31 | 1.34 | 1.33 |
| Clashscore | 5.61 | 5.70 | 6.14 | 4.67 |
| Poor rotamers (%) | 0.78 | 0.52 | 0.00 | 0.52 |
| **Ramachandran plot** |  |  |  |  |
| Favored (%) | 98.83 | 98.25 | 98.45 | 98.25 |
| Allowed (%) | 1.17 | 1.75 | 1.55 | 1.75 |
| Disallowed (%) | 0.00 | 0.00 | 0.00 | 0.00 |

Like previously published OCT and OAT structures^19–21^, URAT1 adopts a major facilitator superfamily (MFS) fold that consists of an extended extracellular domain (ECD), a 12-helical transmembrane domain (TM) and an intracellular helical bundle (ICH). The TM bundle forms a 6+6 pseudosymmetrical arrangement where TMs 1–6 form the N-terminal lobe (N-lobe), and TMs 7–12 comprise the C-terminal lobe (C-lobe).

Interestingly, all the structures we report are of the inward-open conformation, evidenced by the large opening of the central cavity to the intracellular side. All the inhibitors occupy the central binding pocket and make extensive interactions with URAT1_CS_, as if inhibitor binding may stabilize inward-facing states (Fig. 1e). This is notable given that the common mechanism of clinical transporter inhibitors is to stabilize outward-facing conformations^19,22–25^. We therefore sought to explore the functional implications of this mode of binding to URAT1.

### URAT1 drugs are non-competitive inhibitors of uric acid uptake

We conducted a series of uptake experiments in HEK293T cells transiently transfected with hURAT1, where [^14^C]-uric acid and inhibitor are introduced outside the cells and their concentrations were varied to establish the mode of inhibition for each of the compounds tested. We predicted that since the inhibitors occupy the central binding pocket, inhibitors stabilizing outward-facing states will exhibit competitive inhibition whereas those stabilizing inward-facing states will exhibit non-competitive inhibition (Fig. 2a). We found that when comparing non-linear fits to the data of competitive versus non-competitive inhibition, the non-competitive models always resulted in far superior fits (Fig. 2b-d, Table 2). The functional data is consistent with our structural observation that these inhibitors stabilize inward-facing states of URAT1.

**Figure 2.**
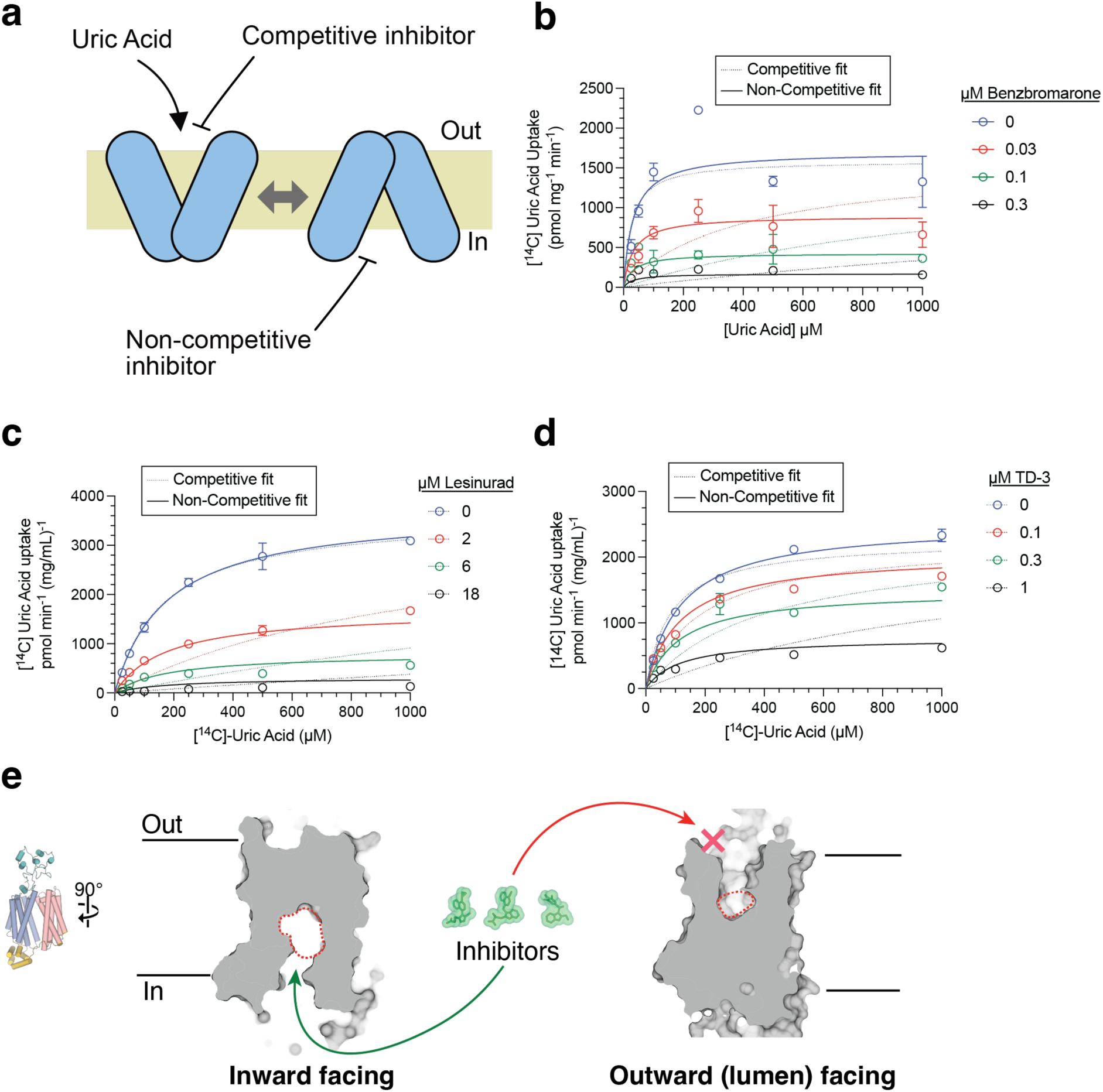
URAT1 inhibitors bind non-competitively to the inward-open conformation. **a,** Schematic of urate uptake by URAT1, and the possible modes of inhibition. **b-d,** Inhibition kinetics determination of [^14^C]-urate uptake (0.9 Ci/mol) for BBR, LESU and TD-3, respectively demonstrating that all three inhibitors inhibit URAT1 non-competitively. Data are presented as mean ± S.E.M (*n* = 3) with global non-linear fits for non-competitive (solid lines) or competitive (dashed lines) models of inhibition. Best fit values and fitting statistics are provided in Table 2. **e,** Comparing the inward-facing (this study) and outward-facing (PDB 8WJQ^26^) URAT1 central cavity size demonstrates the steric restriction for inhibitor binding to the outward facing conformation of URAT1.

**Table 2.** Inhibition kinetics model fitting parameters.

| Inhibitor | Kinetic Model | $K_T$ urate ($\mu\text{M}$)<sup>1</sup> | $K_I$ ($\mu\text{M}$)<sup>1</sup> | $V_{\max}$ ($\text{pmol min}^{-1} \text{mg}^{-1}$)<sup>1</sup> | $Sy.x$<sup>2</sup> |
| --- | --- | --- | --- | --- | --- |
| BBR | *Competitive* | 26.08<br>[11.29 – 48.59] | 0.00213<br>[0.001 – 0.0042] | 1591<br>[1355 – 1846] | 306.7 |
|  | *Non-Competitive* | 32.4<br>[20.7 – 47.9] | 0.033<br>[0.025 – 0.045] | 1700<br>[1530 – 1881] | 231.0 |
| LESU | *Competitive* | 162.8<br>[126.4 – 210.1] | 0.348<br>[0.275 – 0.439] | 3625<br>[3335 – 3955] | 196.3 |
|  | *Non-Competitive* | 175.1<br>[148.8 – 206.5] | 1.63<br>[1.46 – 1.82] | 3725<br>[3522 – 3947] | 137.2 |
| TD-3 | *Competitive* | 82.19<br>[59.19 – 113.0] | 0.079<br>[0.057 – 0.111] | 2259<br>[2071 – 2472] | 208.0 |
|  | *Non-Competitive* | 115.3<br>[99.21 – 130.4] | 0.44<br>[0.38 – 0.50] | 2515<br>[2393 – 2644] | 122.1 |

<sup>1</sup> Fit value with 95% confidence interval [lower value – upper value].
<sup>2</sup> Model fit quality as reported by the standard deviation of the residuals, where
$Sy.x = \sqrt{\frac{\sum(\text{residual}^2)}{n-K}}$ and $n$ is the number of data points (18) and $K$ is the number of fitting parameters (3). When comparing two models, a lower value denotes a better fit. For all inhibitors tested, non-competitive models yield superior fits.

Furthermore, recently reported structures of URAT1 (apo and with uric acid bound) adopt the outward-open conformation^26^. This construct utilized the R477S mutation to stabilize human URAT1 for structural studies, but it also compromises transport activity. Comparing the binding site of the inward- and outward-open conformations of URAT1 reveals that the cavity is far too restrictive in the outward-open conformation to allow inhibitor binding and is much more expansive in the inward-open conformation and (Fig. 2e), explaining why the authors were unable to obtain an inhibitor-bound structure despite their attempts to do so^26^.

The fact that many MFS transporters bind inhibitors in the outward open state is functionally consistent with inhibitors most commonly accessing the transporter from the cell exterior (i.e. blood) to inhibit transport. URAT1 is expressed on the apical membrane in the proximal tubule of kidneys, so URAT1 is exposed to the urine but not to the blood (Fig. 1a). We therefore propose that URAT1 inhibitors bind non-competitively from the intracellular side of the apical membrane (Figs. 1a, 2a). We then wanted to investigate the binding site and probe the functional significance of the residues lining it.

### Central cavity of URAT1

In the URAT1_CS_ structure, the central cavity is mildly conserved (Extended Data Figure 5) and lined with amino acid residues that can be divided into three general groups: a cluster of hydrophobic residues that are distributed on TM2 and TM4 including Y152, L153, I156 and M214, which we termed the hydrophobic region; a cluster of aromatic residues on TM7 and TM5 that spans two opposite sides of the cavity including F241, F360, F364, F365 and F449, which we term the aromatic clamp; and a span of polar or charged residues on TM1, TM4, TM5, and TM8 including S35, T217, N237, S238, D389 and K393 (Fig. 3a). In most MFS-type transporters, TMs 1,4,7 and 10 (termed as A helices) form the central substrate-binding cavity^27^. In contrast, TMs 1, 2, 4, 5, 7, 8 and 10 are all involved in the formation of the central binding cavity of URAT1_CS_ in an inward facing state, indicating that a distinct mechanism might be employed in URAT1 substrate/inhibitor recognition and function.

**Figure 3.**
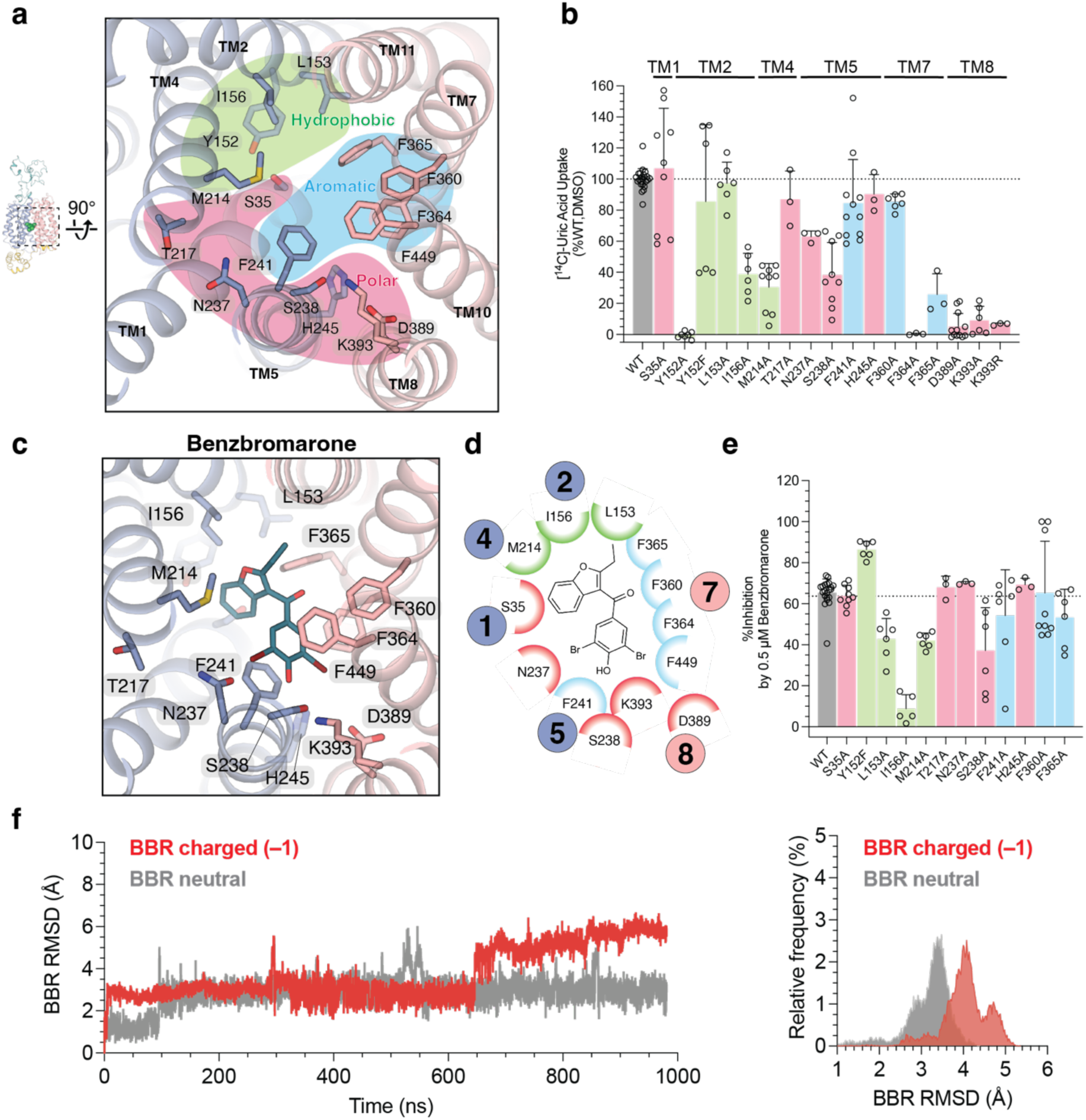
URAT1 central cavity and benzbromarone binding site interactions. **a,** Central cavity of URAT1, using no ligand added URAT1_CS_. **b,** Effects of mutations on central binding cavity residues on uptake of 200 µM [^14^C]-urate (0.9 Ci/mol) in HEK293T cells for 10 min at 37°C in the presence of 1% DMSO. **c,d,** Binding interactions with BBR. Data reported as mean ± standard deviation (S.D.) for *n* = 3-24 replicates **e,** Effects of mutations on inhibition by 0.5 µM BBR on uptake of 200 µM [^14^C]-urate (0.9 Ci/mol) in HEK293T cells for 10 min at 37°C. Data reported as mean ± standard deviation (S.D.) for *n* = 3-21 replicates **f** Left, representative time series trace of root mean squared deviation (R.M.S.D) of charged (red) or neutral (gray) BBR binding in a 1 µs MD simulation. Right, frequency distribution of R.M.S.D. values for charged (red) or neutral (grey) BBR binding over all five replicate MD simulations.

We performed mutagenesis together with radioactive uptake of [^14^C]-uric acid and found that the aromatic and hydrophobic residues on TMs 2 and 7 (Y152, I156, M214, F364, F365) exhibit great effects on uric acid uptake upon mutation (Fig 3b). Notably, F364A abolishes function despite its surface expression (Extended Data Fig. 6). D389 and K393 on TM8 form a salt bridge that is likely more critical to transporter gating than substrate binding directly, as they do not appear close enough to directly interact with uric acid, in agreement with the previous structure^26^. Interestingly, K393 is critical for function, as K393R does not restore activity substantially. Of the critical residues, Y152A is not expressed (Extended Data Figure 6), but Y152F largely restores activity (Fig. 3b).

### Binding of Benzbromarone to URAT1

In our structure of BBR-URAT1_CS_, there is an unambiguous non-protein cryo-EM density centered within the cavity, which allowed us to build the BBR molecule with good confidence and its structure is similar with published BBR structures (Extended Data Figure 7a). BBR forms extensive interactions with the aromatic clamp and occupies the hydrophobic region with its benzofuran group, a position occupied by uric acid in the outward-open conformation^26^ (Extended Data Figure 8). Interestingly, the brominated phenolic group interacts with the aromatic clamp via π-π interaction with F241 and F364. Indeed, the F241A and F365A mutations slightly weaken inhibition by BBR (Fig. 3e). L153A, I156A and M214A, however, have larger effects on inhibition potency, and S238 on TM5 also shows an effect, indicating an important role for these residues for inhibition and a particular importance of the hydrophobic region for BBR binding. To verify the binding mode and stability of BBR binding, molecular dynamics (MD) simulations were conducted on both the charged and neutral forms of BBR, where ionization of the phenolic hydroxyl is readily delocalized across the phenolic ring and extends to the para-carbonyl (Extended Data Figure 7b)^28^. Benzbromarone appears additionally stabilized by interactions of the partially ionized hydroxyl with K393, which is absolutely required for transporter function so its contribution to benzbromarone binding affinity could not be elucidated (Fig. 3b). The MD results in Figure 3f and 3g show the representative R.M.S.D trajectory and histogram for the anionic and neutral forms of BBR within a 1μs timespan, respectively (Extended Data Fig. 9). Neutral BBR, having a lower average R.M.S.D, appears to be more stable inside the cavity compared to the anionic form. This suggests a possible charge interaction with K393 does not significantly contribute to BBR binding and the neutral form of BBR may bind tighter to URAT1.

### Inhibition of URAT1 by lesinurad and TD-3

LESU and TD-3 were modeled confidently into strong, unambiguous densities within the central cavity of URAT1_CS_ (Fig. 1e). For both inhibitors the naphthalene ring (including the bromo/cyclopropyl groups of LESU/TD-3, respectively) largely occupies the hydrophobic region, whereas the heterocycle moieties interact with the aromatic clamp (Fig. 4a, b, d and e). Within the hydrophobic region, M214A has the largest impact on inhibition by LESU (Fig. 4c) and TD-3 (Fig. 4f), in comparison to BBR where I156 plays a more significant role in binding (Fig. 3e). M214 interacts broadly with LESU and TD-3 and specifically with the naphthalene rings of both through a S-π interaction, which is known to impart significant binding stabilization^29^. Unlike BBR, LESU and TD-3 contain mono-carboxylates – localized anions – like the endogenous counter substrates of URAT1^11^. However, while K393 appears to electrostatically stabilize BBR binding, the carboxylates of LESU and TD-3 bind away from K393, appearing instead to potentially hydrogen bond with N237. Mutation of N237 to alanine does not, however, appreciably impact inhibition potency (Fig. 4c, f). M214 also engages with the carboxylate arms of LESU and TD-3. Our MD simulations show stable binding of both drugs (Fig 4g,h) regardless of charge state (Extended Data Fig. 9), but TD-3 shows less mobility within the cavity compared to LESU, in accordance with its stronger binding affinity. Specifically, the carboxylates of both LESU and TD-3 show considerable rotatability during MD simulations, with the carboxyl and dimethyl groups of the carboxylate arm of TD-3 appearing to always interact with M214. A residue that again demonstrates its importance is S238 on TM5, which reduces inhibition potency of not only BBR, but also LESU and TD-3. A picture therefore emerges that rather than highly specific salt bridge interactions between URAT1 and its inhibitors, there is a structural and hydrophobic complementarity with 1-1 interactions provided by the aromatic clamp, S-π interactions from M214, and potential water mediated interactions with S238 on TM5. Notably, based on the structure of urate-bound URAT1, urate overlaps perpendicularly with the location of naphthalene ring of the inhibitors (Extended Data Figure 8). The additional heterocycle and carboxylate of the inhibitors to their respective sites are critical for high affinity binding. Therefore, the interactions mediated by the aromatic clamp and the polar group (both involving TM5) are important, which is consistent with the fact that F241A in TM5 has more impact on LES and TD-3 binding.

**Figure 4.**
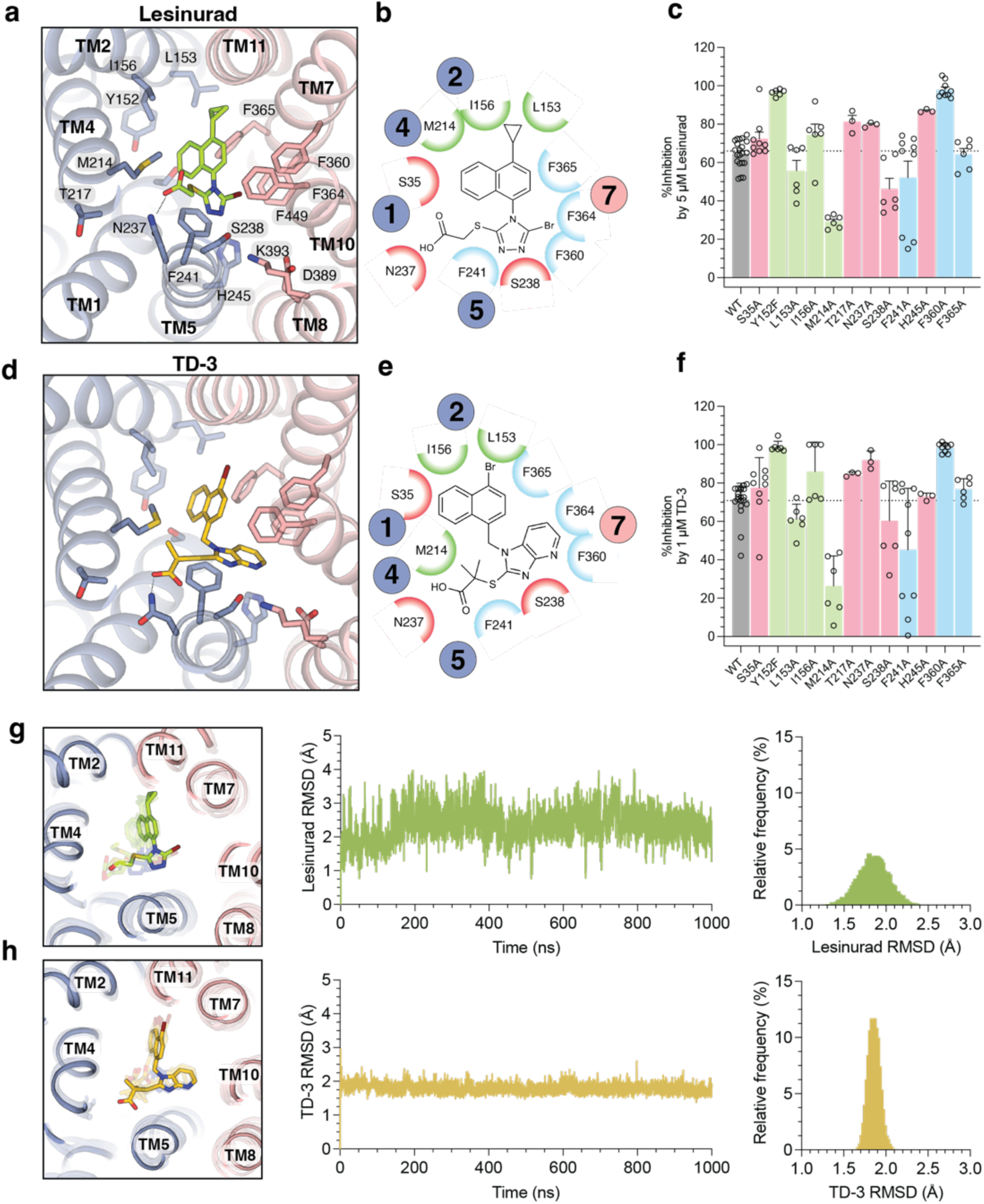
Lesinurad and TD-3 binding site interactions. **a,b,** Binding interactions with LESU. **c,** Effects of mutations on inhibition by 5 µM LESU on uptake of 200 µM [^14^C]-urate (0.9 Ci/mol) in HEK293T cells for 10 min at 37°C. Data reported as mean ± standard deviation (S.D.) for *n* = 3-21 replicates **d,e,** Binding interactions with TD-3. **f,** Effects of mutations on inhibition by 1 µM TD-3 on uptake of 200 µM [^14^C]-urate (0.9 Ci/mol) in HEK293T cells for 10 min at 37°C. Data reported as mean ± standard deviation (S.D.) for *n* = 3-21 replicates. **g,h**, Left, Comparison of cryo-EM structure (no transparency) and MD simulation snapshots (with transparency) of anionic LESU and TD-3 binding to URAT1. Middle, representative R.M.S.D time series trace of LESU and TD-3 binding in 1 µs MD simulations. Right, frequency distribution of R.M.S.D. values for LESU and TD-3 binding, respectively, over all five replicate MD simulations.

### Conformational flexibilities upon inhibitor binding

Despite all our URAT1_CS_ structures being inward-open, directly overlaying the models reveals an ∼10° bend in TM5 of the TD-3-URAT1_CS_ structure, relative to the LESU-and BBR-URAT1_CS_ structures (Fig. 5A). TM5 of URAT1_CS_ alone adopt a conformation similar to that of TD-3 bound URAT1_CS_, likely due to the endogenous molecule bound to the URAT1_CS_ in the absence of inhibitors (Extended Data Fig. 4a). This bend in TM5 originates at G240, in proximity to the previously mentioned S238 residue that is important for inhibitor binding. Importantly, this conformational change is required to accommodate TD-3, where a clash between TD-3 and N237 occurs with the LESU-bound conformation. This observation suggests that there is a conformational ensemble defined by the position of TM5, which can determine inhibitor specificity. Furthermore, unlike for other organic anion/cation transporters, there is no direct specific interaction of the charged substrate/drug moiety with a complementary charged residue^19,20^. While R477 may have a role, the distance between the guanidinium and the charged moieties of these inhibitors are >9Å. The other basic residue, K393 interacts with the phenolic oxygen of BBR, but is ≥8 Å from the carboxylates of LESU and TD-3. A view of the electrostatics of the URAT1_CS_ cavity shows, however, that the region to which these carboxylate moieties or the phenolic ring of BBR occupy is generally electropositive (Fig. 5b). Interestingly, the subtle conformation shift of TM5 in the TD-3 structure induces an electrostatic change in the upper portion of the cavity, which also appears to open slightly larger for solvent access, suggesting that the conformational difference is not limited to TM5 rotation.

**Figure 5.**
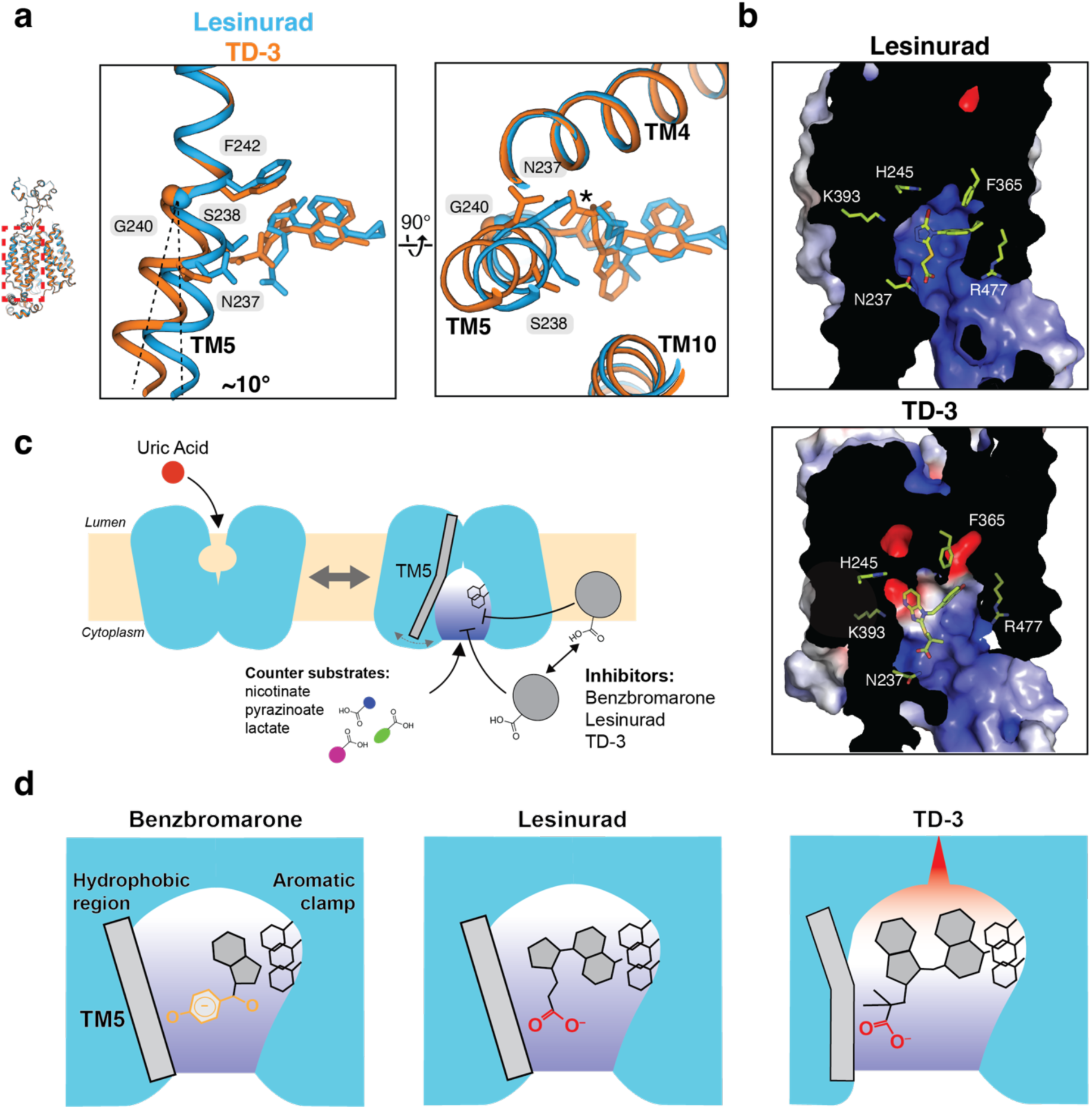
TM5 mobility and binding model for URAT1 inhibitors. **a,** Conformational changes between Les-URAT1_CS_ and TD-3-URAT1_CS_, highlighting TM5 and relevant residues. Note the potential steric clash (*) between lesinurad and N237 in TD-3-URAT1_CS_. **b,** Electrostatic potential surface in Les-URAT1_CS_ (top) and TD-3-URAT1_CS_ (bottom), respectively. **c,** Proposed model for URAT1 substrate transport and inhibition. **d,** Proposed mode for differential inhibition potency among BBR, LESU and TD-3.

## Discussion

Taken together, utilizing cryo-EM, functional studies and molecular dynamics simulations, we have elucidated the inhibitory mechanism of URAT1 by three clinically relevant inhibitors, revealing critical details about their binding poses and the conformational changes upon binding, as summarized in Fig. 5c and 5d. URAT1 is a specific transporter for uric acid, but in exchange transports a variety of mono-carboxylates which have a defined negative charge but vary in size^11^. URAT1, in the outward-open conformation forms a small pocket complementary to uric acid binding from the kidney lumen. Upon changing conformation to the inward-open state, the binding pocket expands into a large electropositive cavity, expelling uric acid and allowing counter substrate binding. This also poses an excellent opportunity for inhibitors to bind to the large, hydrophobic and electropositive cavity of the inward-open URAT1, giving rise to a rather unique non-competitive mode of inhibition. Most inhibitory drugs that target transporters, particularly MFS transporters, lock or stabilize the outward-facing conformation^19,22–25^. Several inhibitor drugs have been found to bind to inward-facing conformations, but this is mostly a feature of the neurotransmitter/sodium symporter family of transporters^23,24,30^. Our data suggest that most URAT1 inhibitors, if not all, likely target the inward-facing states of URAT1. Consistent with this idea, most URAT1 inhibitors are hydrophobic anions which can partition into and pass through the basolateral side of the membrane from the blood, gaining access to URAT1. We posit that this is the optimal strategy for inhibiting not only URAT1, but also other uptake transporters located on the luminal face of the epithelium, like in the gut and kidney.

The variability observed in drug-bound TM5 conformation suggests that multiple sub-conformations of the inward-open state are possible, which may provide greater flexibility in accommodating various anionic counter substrates. This is particularly valuable considering that, without this subtle conformational change, TD-3 cannot bind to URAT1 and that this change drastically modifies the upper cavity electrostatics, opening novel sites for inhibitor interaction. It is unclear whether an induced-fit or conformation selective mechanism is employed in inhibitor binding to URAT1. Given that variously sized monocarboxylates act as counter anions, and that many natural URAT1 inhibitors exist – including multicyclic terpenes and long chain poly-unsaturated fatty acids, which range significantly in size ^13^ – we posit that inhibitors bind to URAT1 via the conformation selection mechanism. The energetic penalty for switching to an inhibitor specific conformation would therefore play a role in inhibitor specificity^31,32^. This feature can be leveraged to achieve greater specificity and efficacy in transporter-targeted drug design.

Our structural, computational, and functional analyses reveal features critical for inhibitor binding. We found that the interactions of the heterocycle and carboxylate groups of the inhibitors with the aromatic clamp and the polar group (both involving TM5) are particularly important. The stronger interactions at these regions make TD-3 a higher affinity inhibitor than LESU. Therefore, further structure-guided optimization of these interactions will be crucial in developing the next generation of URAT1 inhibitors.

Our findings also suggest that the hydrophobic nature of URAT1 inhibitors not only facilitate interactions with the hydrophobic region of the cavity but also increase their effective local concentrations by partitioning into the membrane^33^, contributing to their apparent affinities. BBR has the greatest apparent affinity and in the neutral form has the highest predicted partition coefficient (XLogP3 = 5.7), whereas LESU is less hydrophobic (XlogP3 = 4.7)^34^ and appears to bind less tightly. This difference is expected to be exacerbated for the charged states, where the negative charge on BBR is distributed over the entire phenolic system and carbonyl oxygen (Extended Data Fig. 7b) but concentrated on the carboxylate of LESU and TD-3. High hydrophobicity of BBR would increase its effective concentration substantially, whereas anionic LESU does less well. Consistent with this idea, the MD simulations of BBR binding suggest that direct interactions between BBR and URAT1 are weaker than those of LESU and TD-3. The high hydrophobicity and delocalized negative charge make BBR likely to interact with many off-target membrane proteins, as already reported in its effects on many different classes of membrane and soluble proteins^35–41^. TD-3 has a moderate partitioning but stronger interactions with URAT1 compared to LESU, which results in superior pharmacology, suggesting that a tuning of compound hydrophobicity is required for optimal drug targeting. These differences in charge density and binding may also contribute to drug specificity, as LESU and TD-3 are able to bind with their carboxylates more deeply into the electropositive portion of the cavity. Tailoring carboxylate positioning to perhaps better engage K393 and/or R477 could also be considered for future therapeutic development.

The rising global incidence and suffering caused by gout and hyperuricemia, and the increasing burden on public health systems, necessitates the development of novel inhibitors of URAT1 that exploit the features outlined above. We believe the insights provided by our studies can help achieve more optimal drugs to combat this growing issue.

## Acknowledgements

Cryo-EM data were screened and collected at National Institute of Environmental Health Sciences cryo-EM facility and Duke University Shared Materials Instrumentation Facility, Pacific National Cryo-EM Center (PNCC) and National Cryo-EM Facility (NCEF). We thank Nilakshee Bhattacharya at SMIF, Janette Myers at PNCC and Tara Fox at NCEF for assistance with the microscope operation. This research was supported by Duke Science Technology Scholar Funds (S.-Y.L). We thank Nicholas Wright for valuable advice on this project and Zhenning Ren for critical reading of the manuscript. A portion of this research was supported by the National Institute of Health Intramural Research Program; US National Institutes of Environmental Health Sciences (ZIC ES103326 to M.J.B), the National Cancer Institute’s National Cryo-EM Facility at the Frederick National Laboratory for Cancer Research under contract 75N91019D00024, by the NIH grant U24GM129547 and performed at the PNCC at OHSU and accessed through EMSL (grid.436923.9), a DOE Office of Science User Facility sponsored by the Office of Biological and Environmental Research. DUKE SMIF is affiliated with the North Carolina Research Triangle Nanotechnology Network, which is in part supported by the NSF (ECCS-2025064).

## Author Contributions

Y.S. and J.G.F conducted all single-particle 3D cryo-EM reconstruction and biochemical experiments. H.Z. and S.K. carried out all MD simulations under the guidance of W.I. X.S. synthesized TD-3 under the guidance of P.Z. K.S. screened and collected part of the data under the guidance of M.B. S.-Y.L. and Y.S. performed model building and refinement. K.T. did preliminary biochemical experiments. Y.S. J.G.F. and S.-Y.L. wrote the paper.

## Materials and Methods

### Consensus mutagenesis design

Consensus constructs were designed in a similar manner to what has been previously reported^19,25^, with minor modifications. First, PSI-BLAST was performed to identify 250 hits from UniProt database using human wild-type URAT1 (UniProt ID Q96S37) as query. The hits were manually curated to remove non-URAT1 or incomplete sequences. The remaining sequences were subjected to sequence alignment using MAFFT^42^. The consensus sequence was then extracted in JalView^43^ and aligned to the WT sequence in MAFFT. The final construct features sequence registers consistent with WT.

### HEK293T radiotracer uptake assays

HEK293T cells (ATCC) were cultured in DMEM media (Gibco) supplemented with 10% (v/v) FBS (Gibco) and penicillin-streptomycin (Gibco). The full-length human URAT1 or URAT_CS_ sequences were codon-optimized for *Homo sapiens* and cloned into the BacMam vector with a prescission protease-cleavable C-terminal green fluorescent protein (mEGFP) and FLAG-10xHis purification tags. Site-directed mutagenesis was used to introduce mutations into this background. Empty vector controls utilize the BacMam vector bearing only a FLAG-10xHis-tagged mEGFP. Cells were grown to 60-80% confluency in 10 cm dishes and transfected using 7 µg plasmid DNA and 7 µL TransIT-Pro reagent (Mirus Bio). The next day, cells were detached and transferred to poly-L-lysine treated 24-well plates. After an additional two days at 37°C, the wells were rinsed three times with uptake buffer (25 mM MES-NaOH (pH 5.5), 125 mM Na^+^-gluconate, 4.8 mM K^+^-gluconate, 1.2 mM MgSO_4_, 1.2 mM KH_2_PO_4_, 5.6 mM glucose, 1.3 mM Ca^2+^-gluconate)^44^ and incubated at 37°C for ≥15 min. Uptake was initiated by replacing the media with pre-warmed uptake buffer containing the respective concentrations of [^14^C]-uric acid (Moravek) and inhibitors. Uptake was quenched by addition of ice-cold DPBS (+Ca^2+^/Mg^2+^) then washed thrice by ice-cold DPBS (+Ca^2+^/Mg^2+^). Cells were lysed in 0.1 M NaOH, the protein concentration determined by bicinchoninic acid (BCA) assay, and then transferred to scintillation vials containing EcoLume^TM^ (MP Biomedicals) for counting.

For inhibition kinetics studies, data was fit using GraphPad Prism using competitive (Equation 1) or non-competitive (Equation 2) fitting models^45^, where *K*_T_ is the transport equivalent of the Michaelis constant (*K*_M_), *V*_max_ is the maximal rate of transport, and *K*_I_ is the equilibrium constant for inhibitor binding.

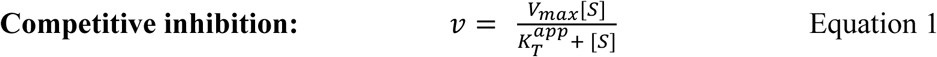

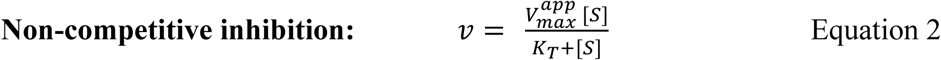

Where 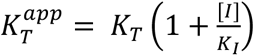 and 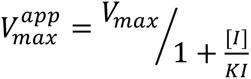

### Surface expression characterization of hURAT1 variants and URAT1_CS_

Surface biotinylation was conducted in 6-well plates on HEK293T cells transiently transfected with the same constructs used for uptake assays, as previously described with modifications^46^. Cells were washed 3x with 1 mL DPBS (+Ca^2+^/Mg^2+^) (Gibco) then incubated for 30 min at 4°C with DPBS (+Ca^2+^/Mg^2+^) containing 0.5 mg mL^-1^ EZ-link Sulfo-NHS-SS-biotin (Thermo Scientific). Biotinylation was quenched by aspirating the biotinylation solution and incubating twice for 5 min with DPBS (+Ca^2+^/Mg^2+^) +100 mM glycine then briefly with unsupplemented DPBS (+Ca^2+^/Mg^2+^). Cells were lysed by addition of 750 µL lysis buffer (20 mM DDM, 50 mM Tris-HCl (pH 8.0), 150 mM NaCl, 10 µg mL^-1^ each of aprotinin, leupeptin and pepstatin, 2 mg mL^-1^ iodoacetamide, and 0.2 mM PMSF) and the lysates transferred to microcentrifuge tubes and incubated for 1h at 4°C. Clarified lysates were quantified by BCA then a consistent amount of total protein across samples was supplemented with additional protease inhibitors and 5 mM EDTA then incubated overnight with 50 µL Neutravidin high-capacity resin slurry (Pierce) at 4°C. The resin was then washed thrice with wash buffer (1 mM DDM, 50 mM Tris-HCl (pH 8.0), 550 mM NaCl) and bound protein eluted with 35 µL of 2x SDS-PAGE sample buffer (BioRad) containing 100 mM dithiothreitol. Following SDS-PAGE (Genscript), protein was transferred onto 0.45 µm PVDF membranes, blocked with 5% bovine serum albumin in Tris-buffered saline and probed with monoclonal mouse anti-FLAG M2 antibody (Sigma Aldrich) diluted 1000x in Tris-buffered saline with 0.1% Tween-20 (TBST), then with IRDye 800CW donkey anti-mouse secondary antibody (LICORbio) diluted 5000x in TBST and imaged with an Odyssey^®^ fluorescence imager (LICORbio).

### URAT1 Protein expression and purification

Full-length consensus URAT1 sequences were codon-optimized for *Homo sapiens* and cloned into the Bacmam vector^60^, in-frame with a PreScission protease cleavage site, followed by EGFP, FLAG-tag and 10× His-tag at the C-terminus. Baculovirus was generated according to manufacturer’s protocol and amplified to P3. For protein expression, HEK293S GnTI^-^ cells (ATCC) was cultured in Freestyle 293 media (Life Technologies) supplemented with 2% (v/v) FBS (Gibco) and 0.5% (v/v) Anti-Anti (Gibco). Cells were infected with 2.5% (v/v) P3 baculovirus at 2.5-3×10^6^ ml^-1^ cell density. After 20 hours shaking incubation at 37°C in the presence of 8% CO_2_, 10 mM sodium butyrate (Sigma-Aldrich) was added to the cell culture and the incubation temperature was lowered to 30°C to boost protein expression. After 40-44 hours, the cells were harvested by centrifugation at 550 × g, and was subsequently resuspended with lysis buffer (20 mM Tris pH 8, 150 mM NaCl, 10 μg mL^-1^ leupeptin, 10 μg mL^-1^ pepstatin, 10 μg mL^-1^ aprotinin, 1 mM phenylmethylsulphonyl fluoride (PMSF, Sigma-Aldrich). The cells were lysed by probe sonication (30 pulses, 3 cycles). The membranes were subsequently solubilized by addition of 1% (w/v) lauryl maltose neopentyl glycol (LMNG, Anatrace), followed by gentle agitation at 4°C for 1 hour. The solubilized lysate was cleared by centrifugation at 16,000 × g for 30 min to remove insoluble material. The supernatant was subsequently incubated with anti-FLAG M2 resin (Sigma-Aldrich) at 4°C for 45 minutes with gentle agitation. The resin was then packed onto a gravity-flow column and washed with 10 column volumes of high-salt wash buffer (20 mM Tris pH 8, 300 mM NaCl, 5mM ATP, 10mM MgSO_4_, 0.005% LMNG), followed by 10 column volumes of wash buffer (20 mM Tris pH 8, 150 mM NaCl, 0.005% LMNG). Protein was then eluted with 5 column volumes of elution buffer (20 mM Tris pH 8, 150 mM NaCl, 0.005% LMNG, 200 μg mL^-1^ FLAG peptide). The eluted protein was concentrated with a 100kDa-cutoff spin concentrator (Millipore), after which 1:10 (w/w) PreScission protease was added to the eluted protein and incubated at 4°C for 1 h to cleave C-terminal tags. The mixture was further purified by injecting onto a Superdex 200 Increase (Cytiva) size-exclusion column equilibrated with GF buffer (20 mM Tris pH 8, 150 mM NaCl, 0.005% LMNG). The peak fractions were pooled and concentrated for cryo-EM sample preparation.

### Cryo-EM sample preparation

The peak fractions from final size exclusion chromatography were concentrated to 9-10 mg ml^-1^. For no ligand added URAT1_CS_ sample, a final concentration of 2% DMSO was added. For ligand added samples (BBR-URAT1_CS,_ LESU-URAT1_CS_, TD-3-URAT1_CS_), 1mM benzbromarone, lesinurad (Sigma-Aldrich) or TD-3 dissolved in DMSO was added 30-40 minutes prior to vitrification. For no ligand added URAT1_CS_ and BBR-URAT1_CS_ samples, protein sample were mixed with a final concentration of 0.5 mM fluorinated octyl maltoside (FOM, Anatrace) prior to vitrification. For les-URAT1_CS_ and TD-3-URAT1_CS_ samples, protein sample were mixed with a final concentration of 0.25 mM FOM prior to vitrification. After mixing with FOM, 3 µL of sample was rapidly applied to a freshly glow-discharged UltrAuFoil R1.2/1.3 300 mesh grids (Quantifoil), blotted with Whatman No. 1 filter paper for 1-1.5 seconds then plunge-frozen in liquid-ethane cooled by liquid nitrogen.

### Cryo-EM data collection

All datasets were collected using a Titan Krios (Thermo Fisher) transmission electron microscope operating at 300 kV equipped with a K3 (Gatan) detector in counting mode behind a BioQuantum GIF energy filter with slit width of 20eV. For no ligand added URAT1_CS_, movies were collected at a nominal magnification of 105,000× with a pixel size of 0.835 Å/px at specimen level, using Latitude S (Gatan) single particle data acquisition program. Each movie was acquired with a nominal dose rate of 19.2 e^-^/px/s over 1.8 s exposure time, resulting a total dose of ∼50 e^-^/Å^2^ over 40 frames. The nominal defocus range was set from −0.7 to –1.7 μm.

BBR-URAT1_CS_ movies were collected at a nominal magnification of 105,000× with a pixel size of 0.855 Å/px at specimen level using Latitude S. Each movie was acquired with a nominal dose rate of 19.3 e^-^/px/s over 2.0 s exposure time, resulting a total dose of ∼50 e^-^/Å^2^ over 40 frames. The nominal defocus range was set from −0.8 to –1.8 μm.

Les-URAT1_CS_ dataset was collected using at a nominal magnification of 105,000× with a super-resolution pixel size of 0.4128 Å/px at specimen level, using SerialEM^47^ data acquisition program. Each movie was acquired with a nominal dose rate of 12.3 e^-^/px/s over 2.0 s exposure time, resulting a total dose of ∼45 e^-^/Å^2^ over 45 frames. The nominal defocus range was set from −1.0 to –2.0 μm.

TD-3-URAT1_CS_ dataset was collected using at a nominal magnification of 105,000× with a pixel size of 0.847 Å/px at specimen level, using SerialEM^47^. Each movie was acquired with a nominal dose rate of 18.2 e^-^/px/s over 2.4 s exposure time, resulting a total dose of ∼60 e^-^/Å^2^ and 60 frames. The nominal defocus range was set from −1.0 to –2.0 μm.

### Cryo-EM data processing

#### No ligand added URAT1_CS_

Beam-induced motion correction and dose-weighing for a total of 18,880 movies were performed using RELION 4.0^48^. Contrast transfer function parameters were estimated using cryoSPARC’s patch CTF estimation^49^. Micrographs showing less than 4.5 Å estimated CTF resolution were discarded, leaving 18,854 micrographs. A subset of 1,500 micrographs were used for blob picking in cryoSPARC^49^, followed by 2D classification to generate templates for template-based particle picking. 2D classes and associated particles that shows the best secondary structure features were used to train a model in Topaz^50^, which were subsequently used for particle picking with Topaz. A total of 6.98 million particles were picked, followed by particle extraction with a 64-pixel box size with 4× binning factor. A reference-free 2D classification was performed to remove obvious junk classes, resulting in a particle set of 6.08 million particles. An iterative ab initio reconstruction triplicate procedure was performed in cryoSPARC, as described previously^19,51^. Four rounds of ab-initio triplicate runs were performed at 4× binned data, resulting in 4.04 million particles. The particles were then re-extracted with 4× binned factor and 6 rounds of ab-initio triplicates were performed, followed by re-extraction without binning factor, at 256-pixel box size wand 2.49 million particles. Twenty-six rounds of ab-initio triplicates were performed with unbinned particle set which resulted in a 527,705 particle set and 3.33 Å resolution reconstruction by non-uniform refinement, and 3.05 Å resolution reconstruction by local refinement with a tight mask covering only protein region. The particle is then transferred to RELION for Bayesian polishing, followed by transferring back to cryoSPARC for local refinement, resulting in a 2.68 Å final reconstruction with 527,705 particles.

### BBR-URAT1_CS_

Benz-URAT1_CS_ dataset was processed similarly to that for no ligand added dataset with minor modifications. Beam-induced motion correction and dose-weighing for a total of 24,488 movies were performed using RELION 4.0^48^. Contrast transfer function parameters were estimated using cryoSPARC’s patch CTF estimation^49^. Micrographs showing less than 4.5 Å estimated CTF resolution were discarded, leaving 21,879 micrographs. A subset of 1,000 images were randomly selected for blob picking, which generated templates for template picking in cryoSPARC, followed by the generation of a 21k particle set for Topaz training. Using Topaz, a 7.73 million particle set was picket. After 2D classification clean-up, 5.50 million particles were retained and subjected to ab-initio triplicate runs. In brief, four, four and 39 rounds of ab-initio triplicate runs were performed at 4× binning, 2× binning and unbinned data sequentially, generating a particle set of 220,530 particles and a 3.29 Å reconstruction by non-uniform refinement. A tight mask covering only protein region was generated using this map and a local refinement using the same particle set and tight mask generated 3.28 Å reconstruction. The particle set were then transferred to RELION for Bayesian polishing, then transferred back to cryoSPARC for non-uniform refinement and local refinements, yielding the final reconstruction of 3.0 Å with 220,530 particles.

### LESU-URAT1_CS_

Les-URAT1_CS_ dataset was processed similarly to that for no ligand added dataset with minor modifications. Beam-induced motion correction and dose-weighing for a total of 13,746 movies were performed using RELION 4.0^48^. During motion correction, the micrographs were two times Fourier binned to generate micrographs with 0.8256 Å/px pixel size. Contrast transfer function parameters were estimated using cryoSPARC’s patch CTF estimation^49^. Micrographs showing less than 4.0 Å estimated CTF resolution were discarded, leaving 13,320 micrographs. A subset of 1,000 images were randomly selected for blob picking, which was used to generate templates for template picking in cryoSPARC, followed by the generation of a 32k particle set for Topaz training. Subsequently, a 9.51 million particle set was picked using trained Topaz model. After two rounds of 2D classification clean-up, 5.04 million particles were retained and subjected to ab-initio triplicate runs. In brief, four, seven and 21 rounds of ab-initio triplicate runs were performed at 4× binning, 2× binning and unbinned data sequentially, yielding a particle set of 512,313 particles and a 3.3 Å reconstruction by non-uniform refinement. The particle set were then transferred to RELION for Bayesian polishing, then transferred back to cryoSPARC for non-uniform refinement and local refinements, with tight mask applied, generating the final reconstruction of 2.74 Å with 512,313 particles.

### TD-3-URAT1_CS_

TD-3-URAT1_CS_ dataset was processed similarly to that for no ligand added dataset with minor modifications. Beam-induced motion correction and dose-weighing for a total of 19,122 movies were performed using RELION 4.0^48^. Contrast transfer function parameters were estimated using cryoSPARC’s patch CTF estimation^49^. Micrographs showing less than 4.5 Å estimated CTF resolution were discarded, leaving 15,790 micrographs. A subset of 500 images were randomly selected for blob picking, which was used to generate templates for template picking in cryoSPARC, followed by the generation of a 56k particle set for Topaz training. Subsequently, a 1.95 million particle set was picked using trained Topaz model. After 2D classification clean-up, 1.65 million particles were retained and subjected to ab-initio triplicate runs. In brief, three and four rounds of ab-initio triplicate runs were performed at 4× binning, 2× binning respectively, yielding a particle set of 1.04 million particles and a 3.3 Å reconstruction by non-uniform refinement. Followed by ab-initio triplicate runs, two rounds of heterogenous refinement was carried out, using three reference classes of the previous obtained 3.3 Å reconstruction without low-pass filtering, low pass filtered to 6 Å and 10 Å, respectively. The class that shows most prominent high resolution features, containing 505,651 particles, was selected and subjected to non-uniform refinement and local refinement with tight masking, yielding a 2.73 Å reconstruction. The particles were then transferred to RELION for Bayesian polishing, then transferred back to cryoSPARC for local refinement, generating a final map of 2.55 Å.

### Model Building and Refinement

All manual model building was performed in Coot^52^ with ideal geometry restraints. A previous OCT1 model (PDB ID 8ET6) was used as an initial reference, followed by further manual model building and adjustment. Idealized CIF restraints for ligands were generated in eLBOW (in PHENIX software suite^53^) from isomeric SMILES strings. After placement, manual adjustments were performed for both protein and ligands ensuring correct stereochemistry and good geometries. The manually refined coordinates were subjected real space refinement in phenix-real.space.refine in PHENIX with global minimization, local grid search and secondary structure restraints. MolProbity^54^ was used to help identify errors and problematic regions. The refined TD-3-URAT1_CS_ cryo-EM structure was then rigid-body fit into the no ligand added URAT1_CS_, BBR-URAT1_CS_ and LESU-URAT1_CS_ maps, followed by manual coordinate adjustments, ligand placement and adjustments, followed by phenix-real.space.refine in PHENIX. The Fourier shell correlation (FSC) of the half- and full-maps against the model, calculated in PHENIX, were in good agreement for all four structures, indicating that the models did not suffer from over-refinement Structural analysis and illustrations were performed using Open Source PyMOL and UCSF Chimera X^55^.

### Molecular Dynamics Simulations

All-atom molecular dynamics (MD) simulations in explicit solvents and POPC bilayer membranes were performed using the cryo-EM BBR-, LESU-, and TD-3-URAT1_CS_ structures. The systems were assembled using CHARMM-GUI *Membrane Builder*.^56–58^ Each system was solvated in TIP3P water and neutralized with 0.15 M Na^+^ and Cl^-^ ions.^59^ Five independent replicates were simulated for each system. Long-range electrostatics in solution were treated with the Particle-mesh Ewald summation,^60,61^ and van der Waals interactions were calculated with a cut-off distance of 9.0 Å. The systems were equilibrated following the CHARMM-GUI *Membrane Builder* protocol. The production runs were performed in the NPT (constant particle number, pressure, and temperature) for 1 μs at 303.15 K and 1 bar with hydrogen mass repartitioning^62,63^ using the following force fields: ff19SB for protein,^64^ OpenFF for ligand, and Lipid21 for lipid.^65^ All simulations were performed with the AMBER22 package^66^ using the system inputs generated by CHARMM-GUI. Ligand binding stability was evaluated by calculating ligand RMSDs after superimposing the TM of the protein structure throughout MD trajectory using CPPTRAJ.^67^

## Extended Data Figures

**Extended Data Figure 1.**
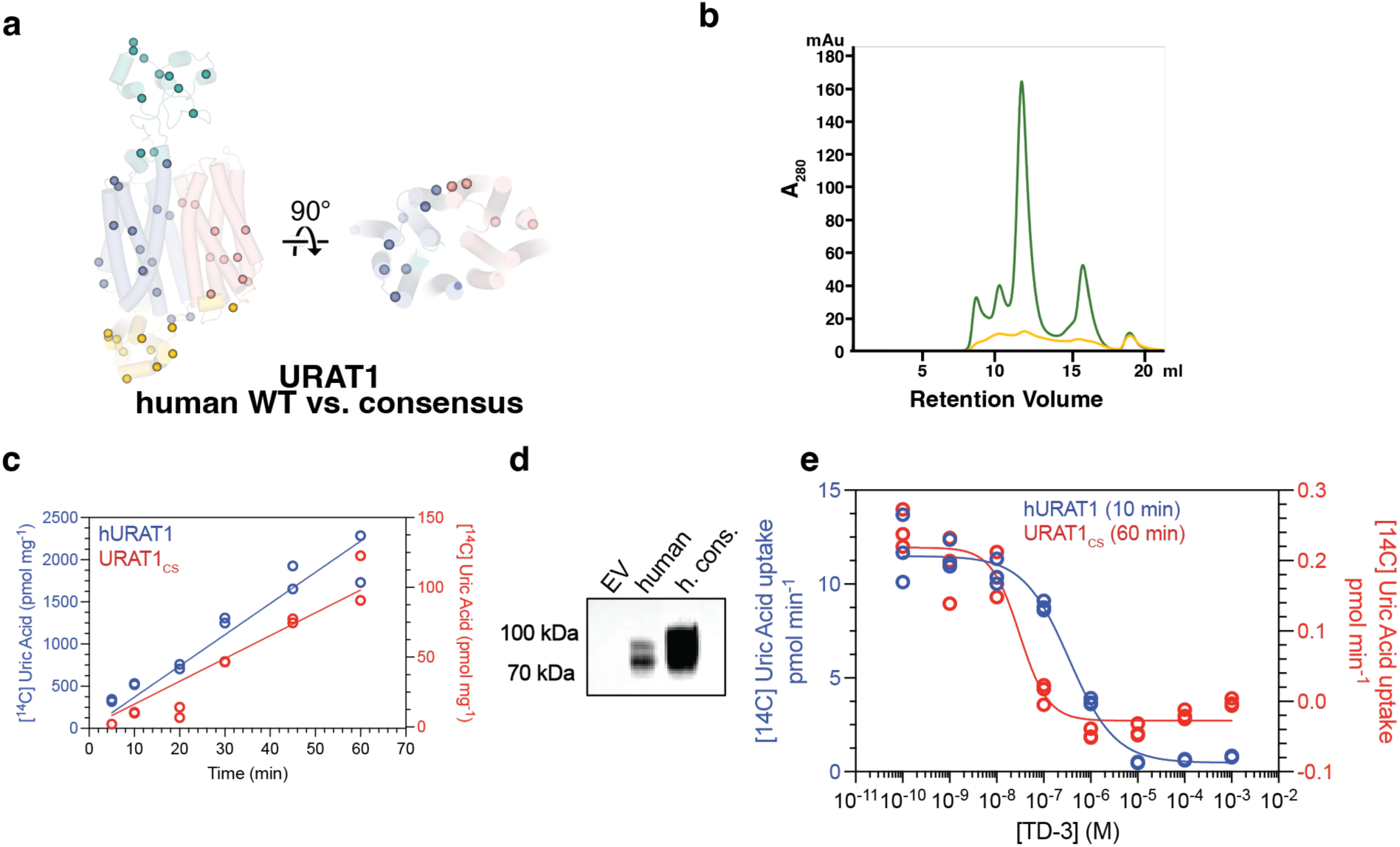
Consensus mutagenesis, functional characterization and protein biochemistry of URAT1_CS_. **a,** Mapping of all the mutations of the consensus construct (URAT1_CS_) relative to the hURAT1 sequence. **b**, Gel filtration profiles of purified hURAT1 (yellow) and URAT1_CS_ (green). **c**, Background-corrected uptake of 10 µM [^14^C]-urate (45 Ci/mol) over time for hURAT1 (left y-axis) and URAT1_CS_ (right y-axis) at 37°C in transiently transfected HEK293T cells. **d,** Surface expression western blot from transiently transfected HEK293T cells showing much greater surface expression of URAT1_CS_ relative to hURAT1. **e**, TD-3 IC_50_ by uptake of 10 µM [^14^C]-urate (45 Ci/mol) in HEK293T cells transiently expressing hURAT1 (10 min at 37°C) or URAT1_CS_ (60 min at 37°C). Background corrected TD-3 titrations were fit to an IC_50_ for hURAT1 of 350 nM [95% CI: 287 – 525 nM], and 31 nM for URAT1_CS_, [95% CI: 17 – 52 nM].

**Extended Data Figure 2.**
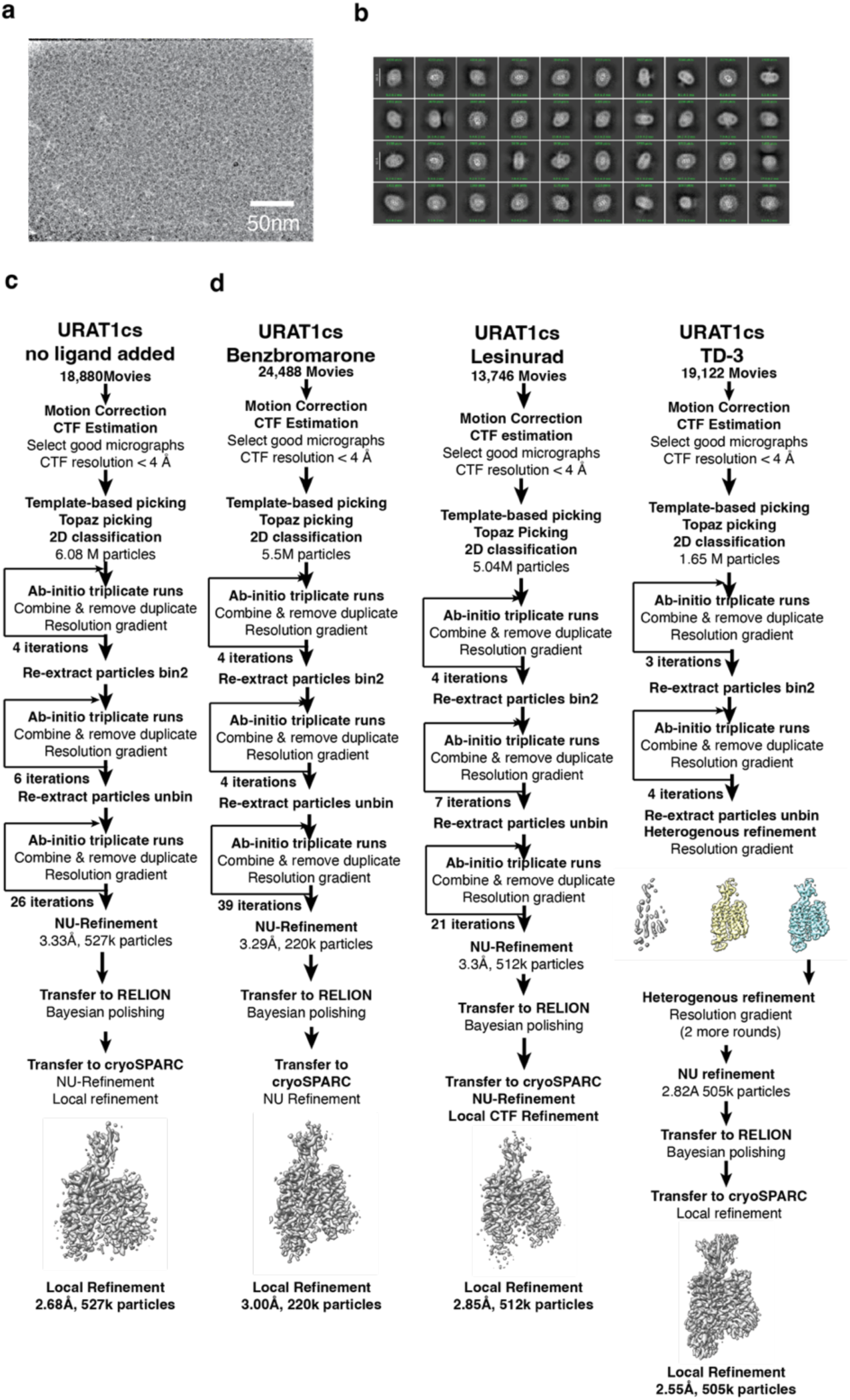
Cryo-EM data processing workflow. Data processing workflow for no ligand added URAT1_CS_, BBR-URAT1_CS_, LESU-URAT1_CS_, and TD-3-URAT1_CS_ datasets, respectively.

**Extended Data Figure 3.**
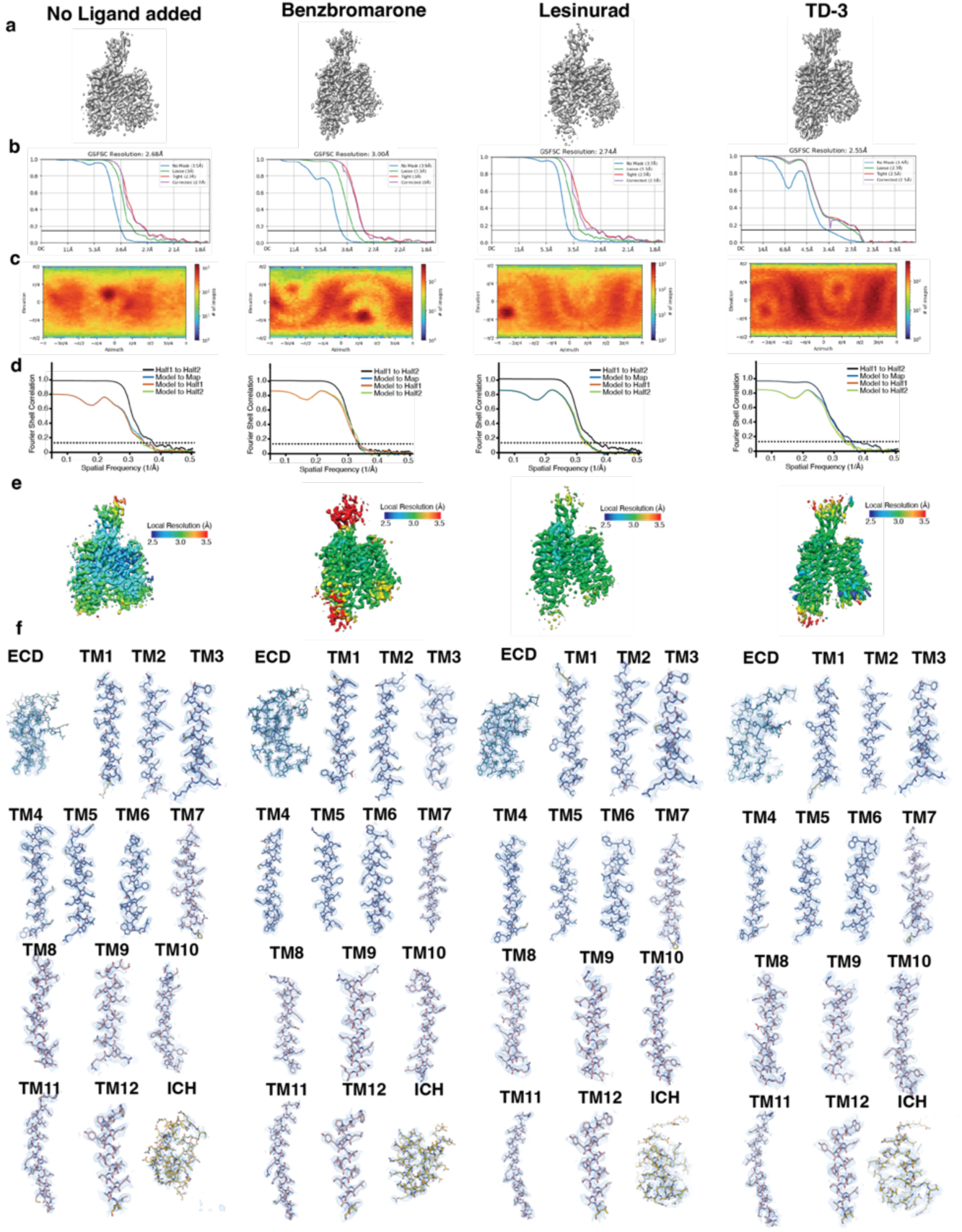
Cryo-EM data validation. **a,** Final cryo-EM reconstructions. **b,** Fourier-shell correlation for the final reconstruction, generated from cryoSPARC. **c,** projection orientation distribution map for the final reconstruction, generated from cryoSPARC. **d,** Map-to-model correlation plots. **e,** Local Resolution plots. **f,** cryo-EM maps for secondary structure segments. From left to right are cryo-EM data validations for URAT1_CS_, BBR-URAT1_CS_, LESU-URAT1_CS_, and TD-3-URAT1_CS_ datasets, respectively.

**Extended Data Figure 4.**
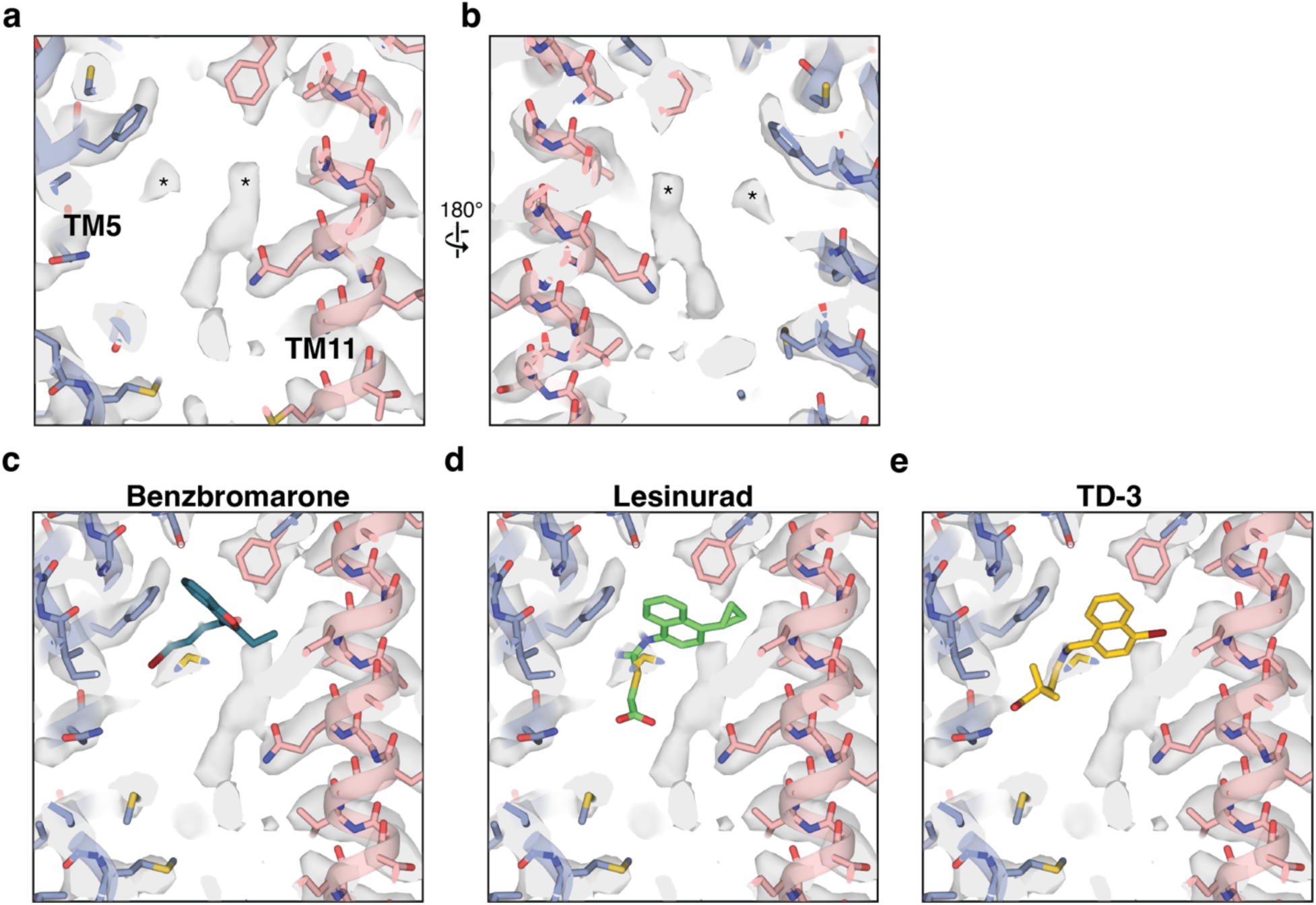
Endogenous cryo-EM peaks in URAT1_CS_ central binding pocket. **a, b,** The appearance of unknown cryo-EM peaks in URAT1_CS_ reconstruction without extra ligand added. **c-e,** the map of URAT1_CS_ overlayed with BBR-URAT1_CS_, LESU-URAT1_CS_, or TD-3-URAT1_CS_ coordinates.

**Extended Data Figure 5.**
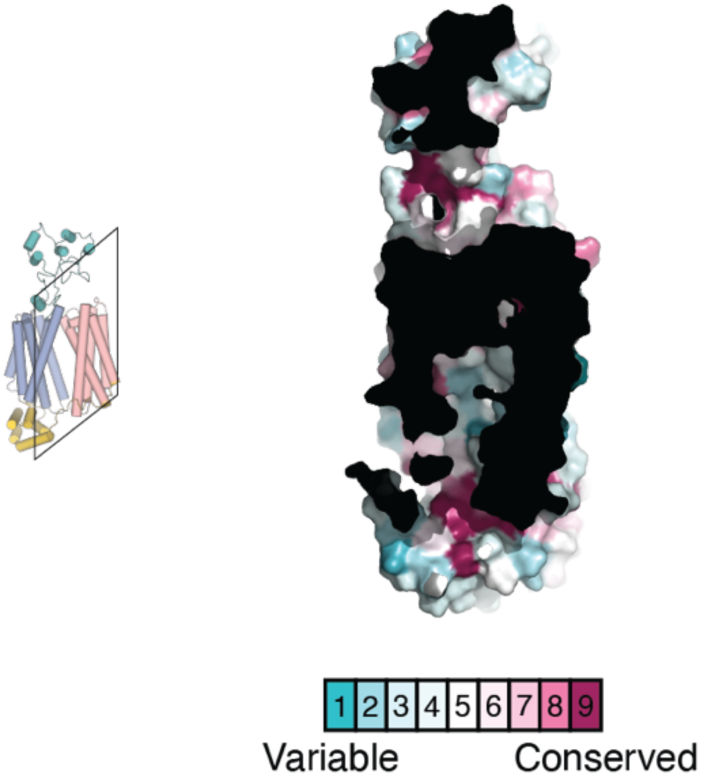
Sequence conservation of URAT1 binding pocket. Consurf analysis^68^ of sequence conservation for URAT1 mapped onto the no inhibitor added structure. The degree of sequence conservation as indicated by the gradient key.

**Extended Data Figure 6.**
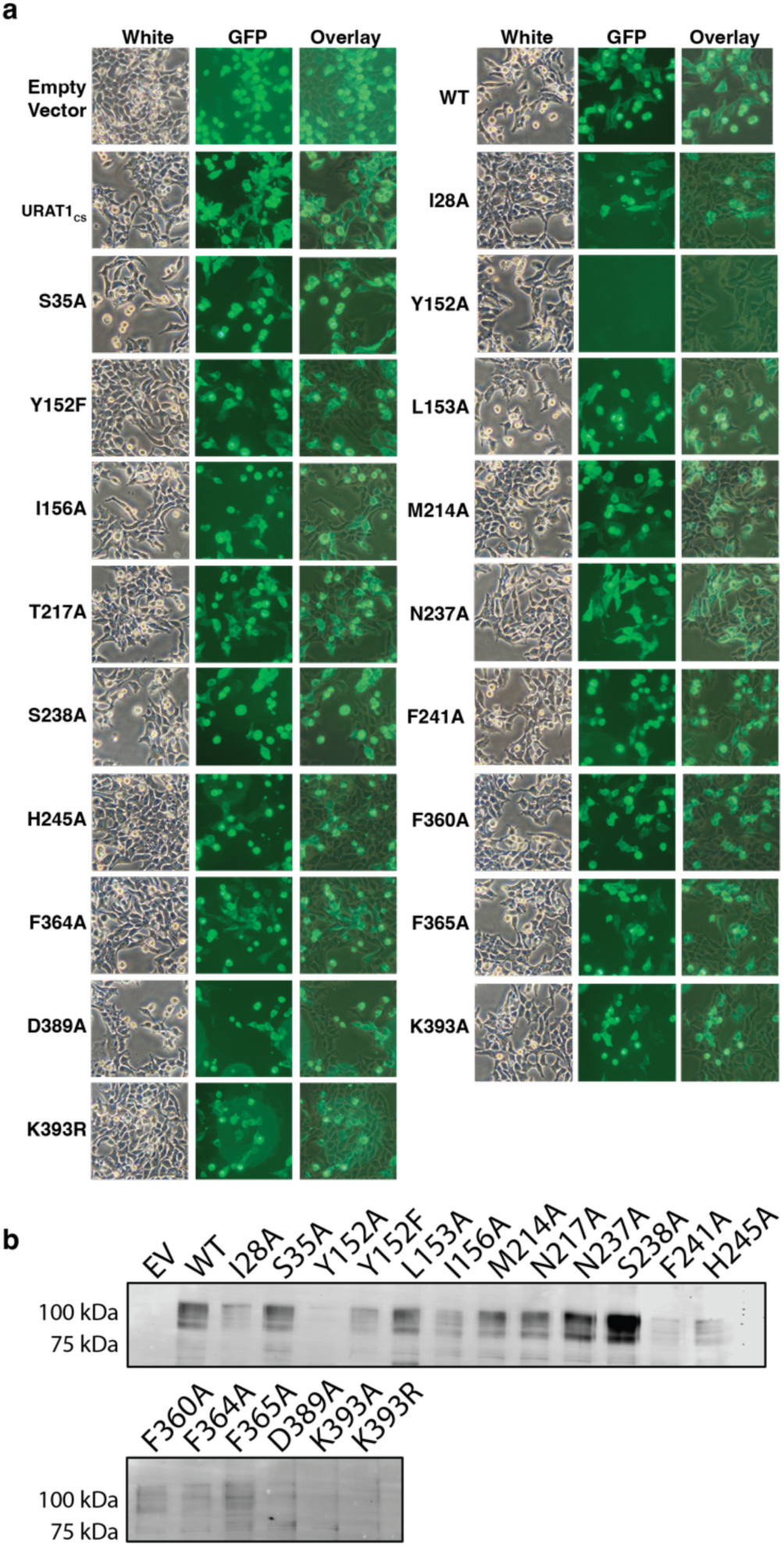
Surface expression of URAT1 and mutants. Microscope images showing bright field, fluorescence and overlay images for the mutants tested in this study. All variants show expression except for Y152A. WT = hURAT1. **b**, Surface expression by surface biotinylation and western blot analysis. EV = empty vector, WT = hURAT1. Only EV and Y152A show no surface expression.

**Extended Data Figure 7.**
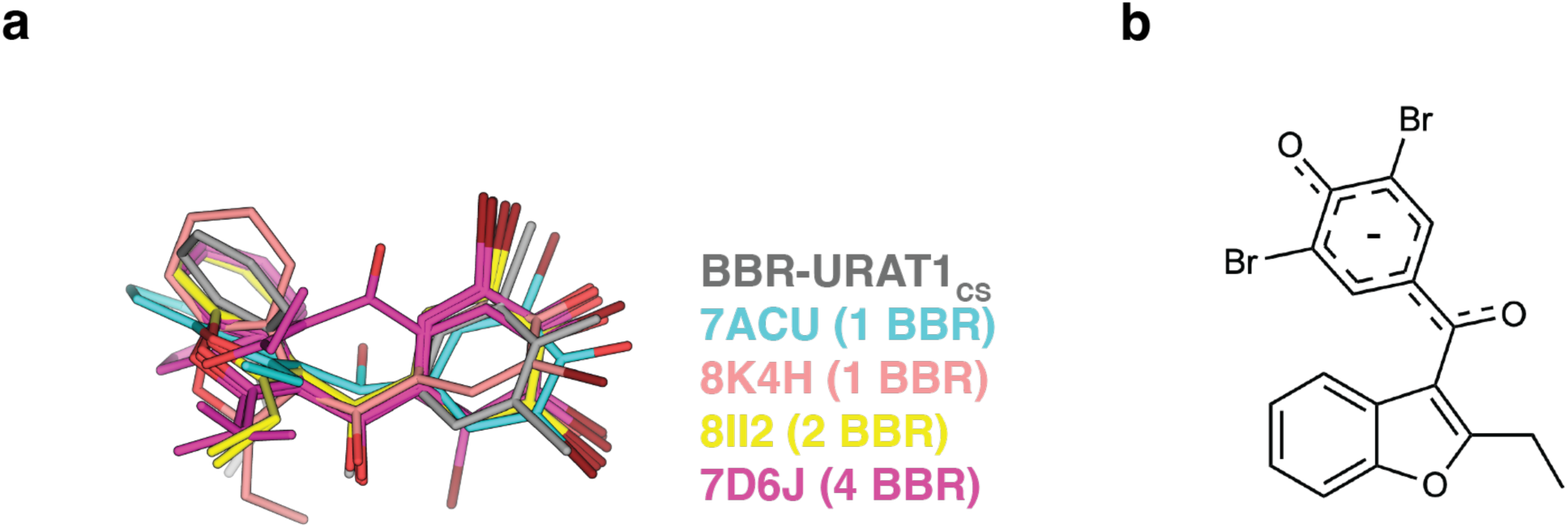
Structural features of BBR. **a,** Overlay of BBR molecule in BBR-URAT1_CS_ with BBR molecules in PDB 7ACU (1 molecule), 8K4H (1 molecule), 8II2 (2 molecules) and 7D6J (4 molecules). BBR conformation in BBR-URAT1_CS_ is similar with 6 out of 8 occurrences. **b,** Resonant charge distribution of BBR at physiological pH, adapted from^28^.

**Extended Data Figure 8.**
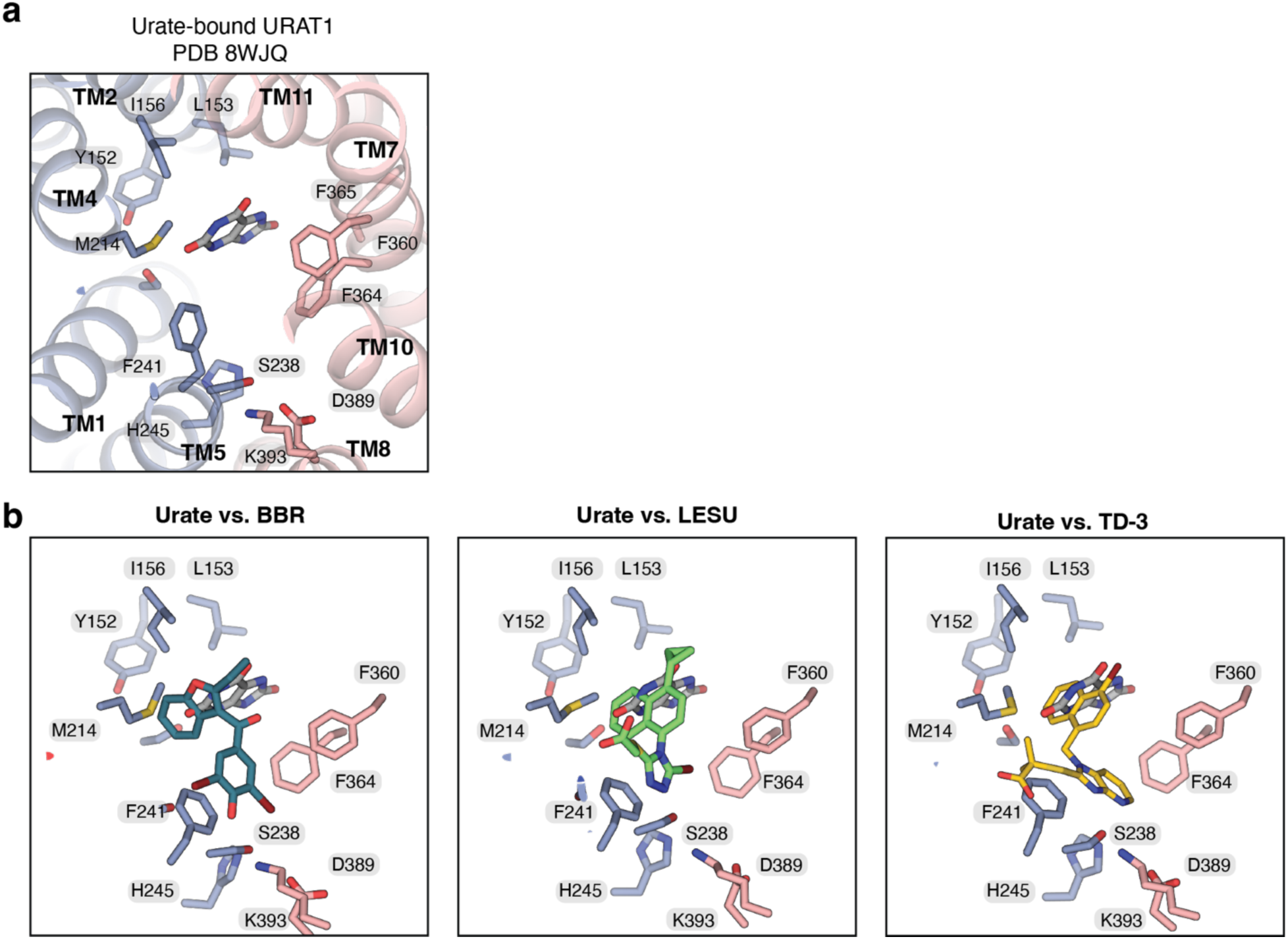
Binding pocket of urate-bound URAT1, adopted from PDB 8WJQ^26^.

**Extended Data Figure 9.**
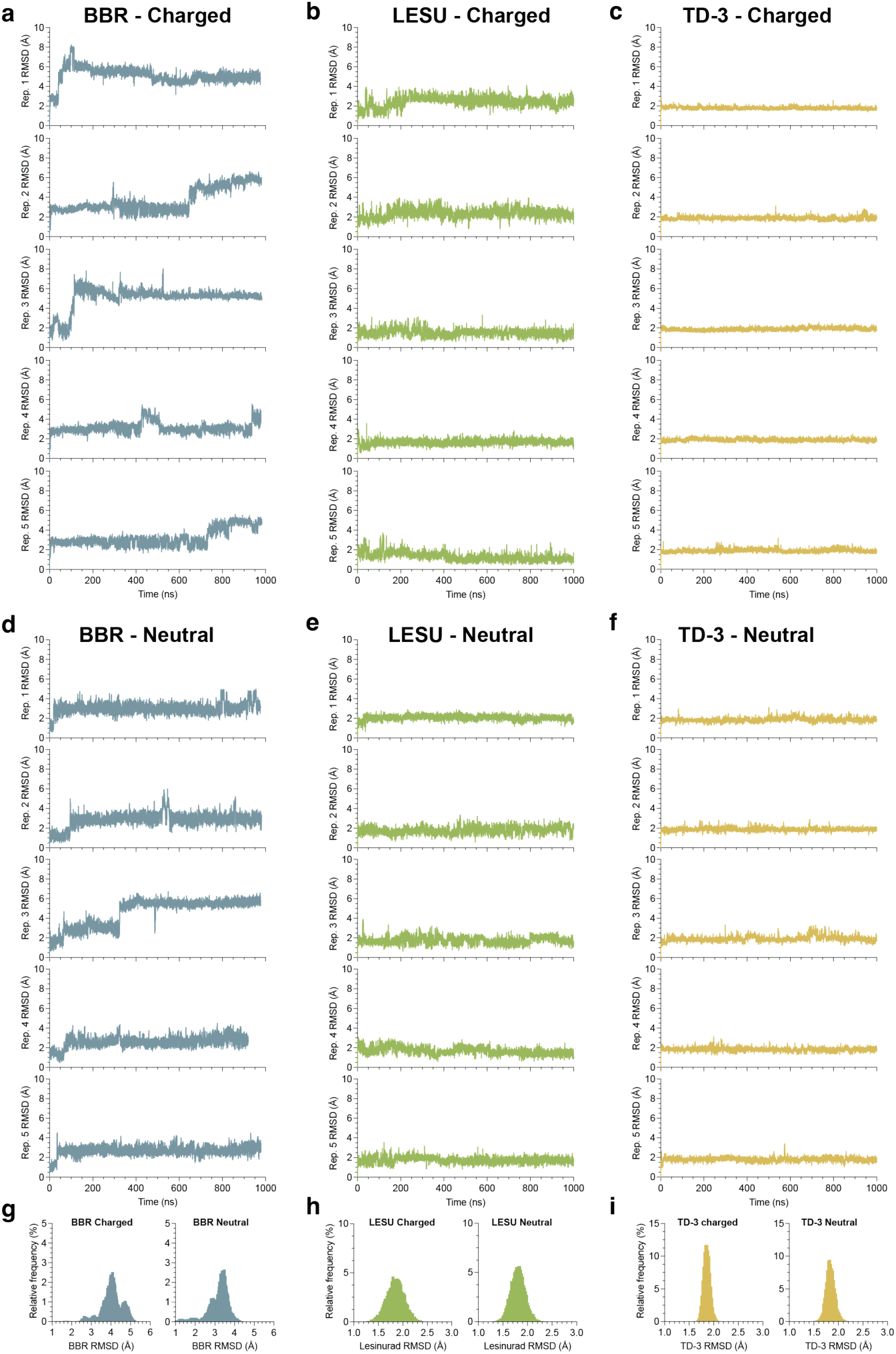
Molecular dynamics for URAT1. Replicate sets of 1 µs simulations for either charged (anionic, –1) or neutral forms of BBR (**a, d**), LESU (**b, e**) and TD-3 (**c, f**). g-i, frequency distribution of RMSD values across all five replicates for charged (left) and neutral (right) forms for BBR (**g**), LESU (**h**) and TD-3 (**i**).

**Supplemental Figure 1.**
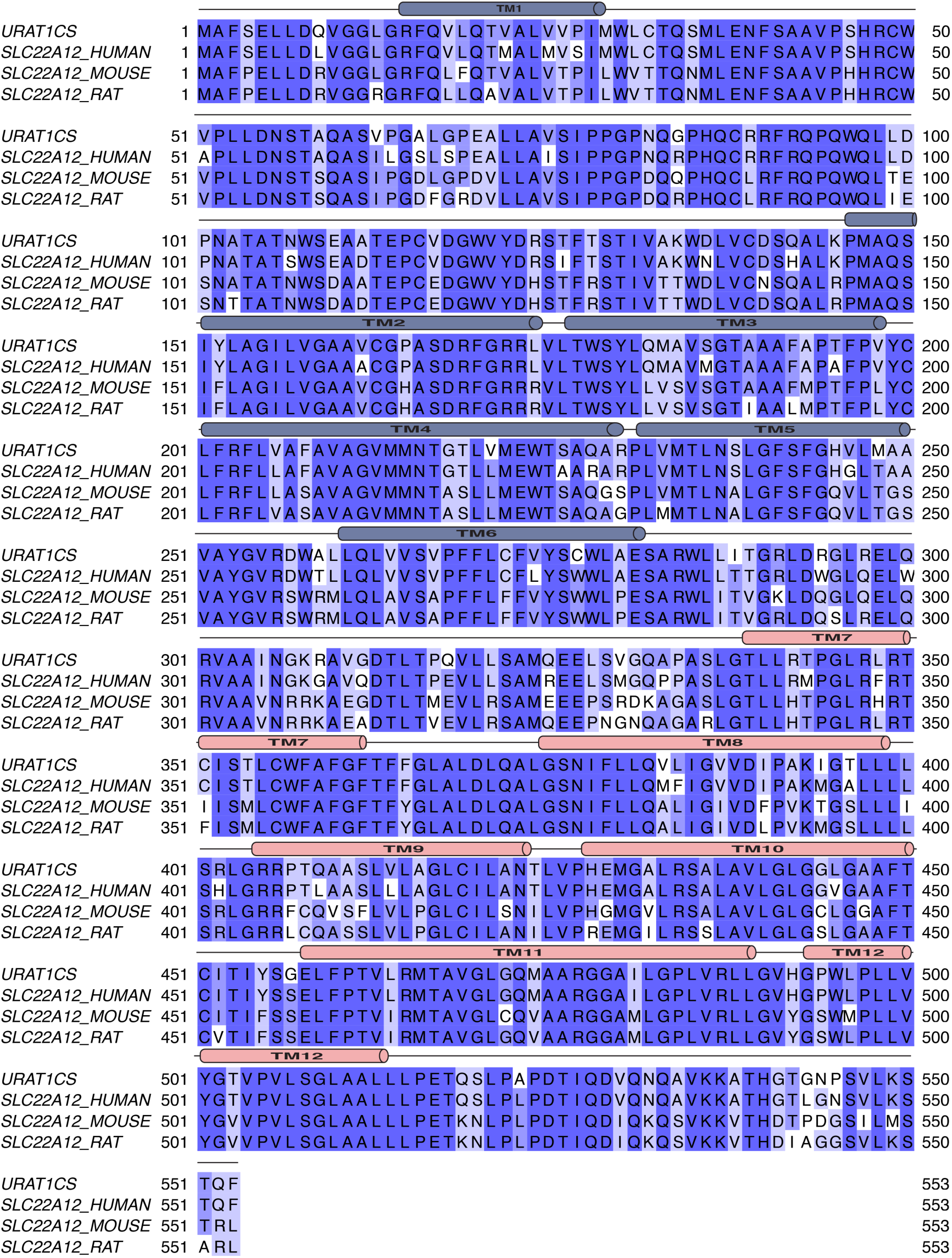
Sequence alignment of URAT1_CS_, human, mouse and rat URAT1 (SLC22A12). Sequence alignment performed using MAFFT^42^.

## References

1 Song, D., Zhao, X., Wang, F. & Wang, G. A brief review of urate transporter 1 (URAT1) inhibitors for the treatment of hyperuricemia and gout: Current therapeutic options and potential applications. European Journal of Pharmacology 907, 174291 (2021).

2 Dehlin, M., Jacobsson, L. & Roddy, E. Global epidemiology of gout: prevalence, incidence, treatment patterns and risk factors. Nature Reviews Rheumatology 16, 380–390 (2020).

3 Punjwani, S. et al. Burden of gout among different WHO regions, 1990–2019: estimates from the global burden of disease study. Scientific Reports 14, 15953 (2024).

4 Skoczyńska, M. et al. Pathophysiology of hyperuricemia and its clinical significance–a narrative review. Reumatologia/Rheumatology 58, 312–323 (2020).

5 Nian, Y.-L. & You, C.-G. Susceptibility genes of hyperuricemia and gout. Hereditas 159, 30 (2022).

6 Du, L. et al. Hyperuricemia and its related diseases: mechanisms and advances in therapy. Signal Transduction and Targeted Therapy 9, 212 (2024).

7 Li, X. et al. Serum uric acid levels and multiple health outcomes: umbrella review of evidence from observational studies, randomised controlled trials, and Mendelian randomisation studies. Bmj 357 (2017).

8 Sharma, G., Dubey, A., Nolkha, N. & Singh, J. A. Hyperuricemia, urate-lowering therapy, and kidney outcomes: a systematic review and meta-analysis. Therapeutic advances in musculoskeletal disease 13, 1759720X211016661 (2021).

9 Yu, Y. et al. Assessing the health risk of hyperuricemia in participants with persistent organic pollutants exposure-a systematic review and meta-analysis. Ecotoxicology and Environmental Safety 251, 114525 (2023).

10 Gu, T. et al. A systematic review and meta-analysis of the hyperuricemia risk from certain metals. Clinical Rheumatology 41, 3641–3660 (2022).

11 Enomoto, A. et al. Molecular identification of a renal urate–anion exchanger that regulates blood urate levels. Nature 417, 447–452 (2002).

12 Xu, L., Shi, Y., Zhuang, S. & Liu, N. Recent advances on uric acid transporters. Oncotarget 8, 100852 (2017).

13 Chen, Y., You, R., Wang, K. & Wang, Y. Recent updates of natural and synthetic URAT1 inhibitors and novel screening methods. Evidence-Based Complementary and Alternative Medicine 2021, 5738900 (2021).

14 Azevedo, V. F., Kos, I. A., Vargas-Santos, A. B., Pinheiro, G. d. R. C. & Paiva, E. d. S. Benzbromarone in the treatment of gout. Advances in Rheumatology 59, 37 (2019).

15 Jansen, T. T. A., Reinders, M., Van Roon, E. & Brouwers, J. Benzbromarone withdrawn from the European market: Another case of” absence of evidence is evidence of absence“?[1]. Clinical and experimental rheumatology 22, 651 (2004).

16 Zhang, M.-Y., Niu, J.-Q., Wen, X.-Y. & Jin, Q.-L. Liver failure associated with benzbromarone: A case report and review of the literature. World Journal of Clinical Cases 7, 1717 (2019).

17 Deeks, E. D. Lesinurad: a review in hyperuricaemia of gout. Drugs & aging 34, 401–410 (2017).

18 Zhao, T. et al. Discovery of novel bicyclic imidazolopyridine-containing human urate transporter 1 inhibitors as hypouricemic drug candidates with improved efficacy and favorable druggability. Journal of Medicinal Chemistry 65, 4218–4237 (2022).

19 Suo, Y. et al. Molecular basis of polyspecific drug and xenobiotic recognition by OCT1 and OCT2. Nature structural & molecular biology 30, 1001–1011 (2023).

20 Dou, T., Lian, T., Shu, S., He, Y. & Jiang, J. The substrate and inhibitor binding mechanism of polyspecific transporter OAT1 revealed by high-resolution cryo-EM. Nature Structural & Molecular Biology 30, 1794–1805 (2023).

21 Parker, J. L., Kato, T., Kuteyi, G., Sitsel, O. & Newstead, S. Molecular basis for selective uptake and elimination of organic anions in the kidney by OAT1. Nature Structural & Molecular Biology 30, 1786–1793 (2023).

22 Niu, Y. et al. Structural basis of inhibition of the human SGLT2–MAP17 glucose transporter. Nature 601, 280–284 (2022).

23 Wu, D. et al. Transport and inhibition mechanisms of human VMAT2. Nature 626, 427–434 (2024).

24 Li, Y. et al. Dopamine reuptake and inhibitory mechanisms in human dopamine transporter. Nature, 1–9 (2024).

25 Wright, N. J. & Lee, S.-Y. Structures of human ENT1 in complex with adenosine reuptake inhibitors. Nature structural & molecular biology 26, 599–606 (2019).

26 He, J., et al. Structural basis for the transport and substrate selection of human urate transporter 1. Cell Reports 43 (2024).

27 Drew, D., North, R. A., Nagarathinam, K. & Tanabe, M. Structures and general transport mechanisms by the major facilitator superfamily (MFS). Chemical reviews 121, 5289–5335 (2021).

28 Locuson, C. W., Suzuki, H., Rettie, A. E. & Jones, J. P. Charge and substituent effects on affinity and metabolism of benzbromarone-based CYP2C19 inhibitors. Journal of medicinal chemistry 47, 6768–6776 (2004).

29 Suo, Y. et al. Structural insights into electrophile irritant sensing by the human TRPA1 channel. Neuron 105, 882–894. e885 (2020).

30 Hu, T. et al. Transport and inhibition mechanisms of the human noradrenaline transporter. Nature, 1–8 (2024).

31 Yin, Y. et al. Mechanisms of sensory adaptation and inhibition of the cold and menthol receptor TRPM8. Science Advances 10, eadp2211 (2024).

32 Afek, A. et al. DNA mismatches reveal conformational penalties in protein–DNA recognition. Nature 587, 291–296 (2020).

33 Lee, S.-Y. & MacKinnon, R. A membrane-access mechanism of ion channel inhibition by voltage sensor toxins from spider venom. Nature 430, 232–235 (2004).

34 Cheng, T. et al. Computation of octanol− water partition coefficients by guiding an additive model with knowledge. Journal of chemical information and modeling 47, 2140–2148 (2007).

35 Gao, J. et al. A novel role of uricosuric agent benzbromarone in BK channel activation and reduction of airway smooth muscle contraction. Molecular Pharmacology 103, 241–254 (2023).

36 Huang, F. et al. Calcium-activated chloride channel TMEM16A modulates mucin secretion and airway smooth muscle contraction. Proceedings of the National Academy of Sciences 109, 16354–16359 (2012).

37 Zheng, Y. et al. Activation of peripheral KCNQ channels relieves gout pain. Pain 156, 1025–1035 (2015).

38 Tang, L. W. T. et al. Differential reversible and irreversible interactions between benzbromarone and human cytochrome P450s 3A4 and 3A5. Molecular Pharmacology 100, 224–236 (2021).

39 Cotrina, E. Y. et al. Repurposing benzbromarone for familial amyloid polyneuropathy: a new transthyretin tetramer stabilizer. International Journal of Molecular Sciences 21, 7166 (2020).

40 Cai, H.-y., et al. Benzbromarone, an old uricosuric drug, inhibits human fatty acid binding protein 4 in vitro and lowers the blood glucose level in db/db mice. Acta Pharmacologica Sinica 34, 1397–1402 (2013).

41 Loo, T. W., Bartlett, M. C. & Clarke, D. M. Benzbromarone stabilizes ΔF508 CFTR at the cell surface. Biochemistry 50, 4393–4395 (2011).

42 Katoh, K. & Toh, H. Recent developments in the MAFFT multiple sequence alignment program. Briefings in bioinformatics 9, 286–298 (2008).

43 Waterhouse, A. M., Procter, J. B., Martin, D. M., Clamp, M. & Barton, G. J. Jalview Version 2—a multiple sequence alignment editor and analysis workbench. Bioinformatics 25, 1189–1191 (2009).

44 Tan, P. K., Ostertag, T. M. & Miner, J. N. Mechanism of high affinity inhibition of the human urate transporter URAT1. Scientific reports 6, 34995 (2016).

45 Cornish-Bowden, A. Fundamentals of enzyme kinetics. (John Wiley & Sons, 2013).

46 Huang, G. N. Biotinylation of cell surface proteins. Bio-protocol 2, e170–e170 (2012).

47 Mastronarde, D. N. SerialEM: a program for automated tilt series acquisition on Tecnai microscopes using prediction of specimen position. Microscopy and Microanalysis 9, 1182–1183 (2003).

48 Zivanov, J. et al. A Bayesian approach to single-particle electron cryo-tomography in RELION-4.0. Elife 11, e83724 (2022).

49 Punjani, A., Rubinstein, J. L., Fleet, D. J. & Brubaker, M. A. cryoSPARC: algorithms for rapid unsupervised cryo-EM structure determination. Nature methods 14, 290–296 (2017).

50 Bepler, T. et al. Positive-unlabeled convolutional neural networks for particle picking in cryo-electron micrographs. Nature methods 16, 1153–1160 (2019).

51 Wright, N. J. et al. Methotrexate recognition by the human reduced folate carrier SLC19A1. Nature 609, 1056–1062 (2022).

52 Emsley, P., Lohkamp, B., Scott, W. G. & Cowtan, K. Features and development of Coot. Acta Crystallographica Section D: Biological Crystallography 66, 486–501 (2010).

53 Liebschner, D. et al. Macromolecular structure determination using X-rays, neutrons and electrons: recent developments in Phenix. Acta Crystallographica Section D: Structural Biology 75, 861–877 (2019).

54 Williams, C. J. et al. MolProbity: more and better reference data for improved all-atom structure validation. Protein Science 27, 293–315 (2018).

55 Meng, E. C. et al. UCSF ChimeraX: Tools for structure building and analysis. Protein Science 32, e4792 (2023).

56 Jo, S., Kim, T., Iyer, V. G. & Im, W. CHARMM-GUI: a web-based graphical user interface for CHARMM. J Comput Chem 29, 1859–1865 (2008). 10.1002/jcc.20945

57 Wu, E. L. et al. CHARMM-GUI Membrane Builder toward realistic biological membrane simulations. J Comput Chem 35, 1997–2004 (2014). 10.1002/jcc.23702

58 Lee, J. et al. CHARMM-GUI Membrane Builder for Complex Biological Membrane Simulations with Glycolipids and Lipoglycans. Journal of Chemical Theory and Computation 15, 775–786 (2019). 10.1021/acs.jctc.8b01066

59 Jorgensen, W. L., Chandrasekhar, J., Madura, J. D., Impey, R. W. & Klein, M. L. Comparison of Simple Potential Functions for Simulating Liquid Water. J Chem Phys 79, 926–935 (1983). 10.1063/1.445869

60 Darden, T., York, D. & Pedersen, L. Particle Mesh Ewald - an N.Log(N) Method for Ewald Sums in Large Systems. J Chem Phys 98, 10089–10092 (1993). 10.1063/1.464397

61 Essmann, U. et al. A Smooth Particle Mesh Ewald Method. J Chem Phys 103, 8577–8593 (1995). 10.1063/1.470117

62 Hopkins, C. W., Le Grand, S., Walker, R. C. & Roitberg, A. E. Long-Time-Step Molecular Dynamics through Hydrogen Mass Repartitioning. Journal of Chemical Theory and Computation 11, 1864–1874 (2015). 10.1021/ct5010406

63 Gao, Y. et al. CHARMM-GUI Supports Hydrogen Mass Repartitioning and Different Protonation States of Phosphates in Lipopolysaccharides. Journal of Chemical Information and Modeling 61, 831–839 (2021). 10.1021/acs.jcim.0c01360

64 Tian, C. et al. ff19SB: Amino-Acid-Specific Protein Backbone Parameters Trained against Quantum Mechanics Energy Surfaces in Solution. J Chem Theory Comput 16, 528–552 (2020). 10.1021/acs.jctc.9b00591

65 Dickson, C. J., Walker, R. C. & Gould, I. R. Lipid21: Complex Lipid Membrane Simulations with AMBER. J Chem Theory Comput 18, 1726–1736 (2022). 10.1021/acs.jctc.1c01217

66 Case, D. A. et al. AmberTools. J Chem Inf Model 63, 6183–6191 (2023). 10.1021/acs.jcim.3c01153

67 Roe, D. R. & Cheatham, T. E., 3rd. PTRAJ and CPPTRAJ: Software for Processing and Analysis of Molecular Dynamics Trajectory Data. J Chem Theory Comput 9, 3084–3095 (2013). 10.1021/ct400341p

68 Yariv, B. et al. Using evolutionary data to make sense of macromolecules with a “face-lifted” ConSurf. Protein Science 32, e4582 (2023).

